# *Synaptopodin* KO rat for assessing the dendritic spine apparatus and axonal cisternal organelle in synaptic plasticity, development, and behavior

**DOI:** 10.1101/2025.11.25.690444

**Authors:** Masaaki Kuwajima, Olga I. Ostrovskaya, Lyndsey M. Kirk, Ashley Alario, Weiling Yin, Sonia Singh, Anna Xaymongkhol, Adrienne Li, Ella Prasad, Kristen M. Harris

## Abstract

The actin-binding protein synaptopodin (Synpo) regulates the cytoskeleton and organization of endoplasmic reticulum, thereby amplifying intracellular Ca^2+^ signaling. Knockout (KO) mouse models have been used to study the role of Synpo in kidney and brain functions, where it supports stress fiber formation, as well as long-term potentiation (LTP) and learning, respectively. Here, we generated *Synpo* KO rats using CRISPR-Cas9, and show they are viable but have reduced body weight after postnatal days 35-45, along with shorter limb bone length. Their basal kidney function is normal into early adulthood. Serial section electron microscopy from *Synpo* KO rat hippocampus reveals the absence of the spine apparatus in dendrites and cisternal organelle in the axon initial segment (AIS), two Synpo-dependent specializations of smooth endoplasmic reticulum. The AIS in KO was still innervated by inhibitory synapses despite the total loss of the cisternal organelle. *Synpo* KO rats also showed reduced LTP. Previously unknown KO effect of Synpo on body stature could have an inadvertent impact on behavioral outcomes. Furthermore, rats have a well-defined developmental onset of LTP, compared to the variable onset of LTP in mice. This, combined with known species differences in behavior, makes our KO rat model a valuable resource for assessing the role of Synpo in development, learning, synaptic plasticity, and a wide range of other biological functions.

## Introduction

Regulation of actin cytoskeleton and intracellular Ca^2+^ is essential for cellular functions, including synaptic plasticity in neurons and glomerular filtration in the kidney. The actin-binding protein synaptopodin (Synpo) is expressed in both neurons and kidney podocytes. Synpo is critically involved in controlling the structure of smooth endoplasmic reticulum (SER) [1–3] that affects Ca^2+^ transients [4]. Synpo regulates actin bundling and stress fiber formation [5], and change in its expression level has been proposed as a biomarker for glomerular diseases, mild cognitive impairment, Alzheimer’s disease, Lewy body dementias, multiple sclerosis, and acute hypoxic brain injury [6–13]. Although *Synpo* transcripts are expressed widely in the body [14–17] (https://www.gtexportal.org/home/gene/SYNPO), most studies thus far focused on the role of Synpo protein in brain and kidney functions.

In the foot processes of podocytes in the kidney glomerulus, the long and T isoforms of Synpo promote actin bundling and stress fiber formation mediated by alpha-actinin [5]. This process is regulated by phosphorylation-dependent proteolysis of Synpo [18–20]. Loss of Synpo compromises the foot process function, impairing recovery from nephrotic syndrome induced by the bacterial component lipopolysaccharides [5] or the chemotherapy drug Adriamycin [21].

The short isoform of Synpo is expressed in many forebrain areas, where it is required for the formation of the spine apparatus in dendritic spines and cisternal organelles in axon initial segments. Both structures are derived from multiple cisterns of SER integrated with the actin cytoskeleton and Synpo [1–3,22,23]. The spine apparatus mediates Ca^2+^-induced Ca^2+^ signaling in dendritic spines [4,24,25] and may serve as a local secretory pathway compartment capable of ribosome docking and post-translational modification of integral membrane proteins, including glycosylation [26–29]. Synpo is upregulated during long-term potentiation (LTP), a widely studied mechanism of learning and memory [14,30]. Spines containing the spine apparatus undergo greater amounts of plasticity than those lacking this organelle [31–33]. The cisternal organelle also serves as a source of intracellular Ca^2+^ and plays a role in plasticity of the axon initial segment and regulation of neuronal excitability [2,34–37].

We previously found that LTP induced by theta-burst stimulation (TBS) in the hippocampal area CA1 of adult rats is associated with the conversion of SER tubules into the spine apparatus, as well as synapse enlargement and clustering around spines containing spine apparatus [33]. In contrast, TBS-induced LTP in juvenile (P15) rats is linked to synaptogenesis rather than SER-associated synapse enlargement [38,39]. Interestingly, Synpo expression is developmentally regulated, with protein expression beginning at P5 and reaching adult levels by P12 [40,41]. Previous work suggests that the spine apparatus is absent at birth and requires maturation past P15 in rat hippocampus [42,43]. Accordingly, the developmental upregulation of Synpo and formation of the spine apparatus may contribute to the maturation of synapses, spine clustering, and LTP [33,39].

A mouse model of *Synpo* deletion has been used to study the role of Synpo in kidney and brain functions. This animal model has revealed that Synpo and the spine apparatus are involved in the induction and maintenance of LTP and synaptic plasticity [1,44–48]. Mice have become the genetic model of choice since the first gene knockout model was generated in 1987 [49]. More recently, advances in gene editing methods, such as approaches based on clustered regularly interspaced short palindromic repeats (CRISPR)-Cas [50], are expanding the utility of rats as genetic models. Rats and mice are often regarded as interchangeable due to their morphological and evolutionary similarity. However, a growing body of evidence demonstrates that their genetic and functional differences need to be considered when choosing the model system best suited for a specific biological question. For example, the onset of LTP expression in the mouse hippocampus area CA1 is highly variable over postnatal day (P) 18-37 [51], while rats have a well-defined onset age for LTP at P12 that coincides with spinogenesis [52,53]. Rats and mice also differ in their navigational strategies and the stability of neural representation of space, even though they share many behavioral characteristics in simple exploration tasks (reviewed in [54]). In addition, rats are more receptive to human handling routines and less easily stressed by human contact than mice [55,56]. These findings point to species differences that should be taken into account for establishing the functional role of Synpo.

Here, we generate and investigate *Synpo* knockout (KO) rats. The outcomes demonstrate that KO rats are an important resource for probing the impact of *Synpo* deletion on brain, bone, and kidney functions, especially during development, learning, memory, and a variety of disease processes.

## Results

We used the CRISPR-Cas9 system to delete the coding region of the *Synpo* gene in rats (Figs 1A and B, S1 Fig), designing guide RNAs to target the 5’ and 3’ regions flanking the protein-coding exons 2 and 3. The successful deletion was confirmed by sequencing the PCR products spanning the excised region from N1 generation rats (S1 Document). The resulting two KO lines carry alleles that differ in the genomic location of the 5’ breakpoint (S1 Document). The KO rats are viable and appear healthy into adulthood under standard laboratory housing conditions. When heterozygotes were paired for breeding, the resulting litter size averaged 11, ranging from 4 to 18 pups (based on 51 litters from 20 breeding pairs). Both KO lines were used to measure body weight (Fig 2C) and were prepared for electron microscopy to ensure deletion of the spine apparatus (Fig 3) and cisternal organelle (Fig 4). After the first line was deposited to the Rat Resource and Research Center (Columbia, MO; https://www.rrrc.us), under the designation LE-Synpo*^em1Kmh^*, the second line (LE-Synpo*^em2Kmh^*) was used for histology (Figs 1D and E), renal function tests (Fig 2A), bone length measurements (Fig 2D), and slice physiology (Fig 5). WT littermates were used as control as much as possible. Detailed information about the animals used (including founder lines and littermates) is available in additional data deposited at the Texas Data Repository (see “Data and code availability”).

**Fig 1.**
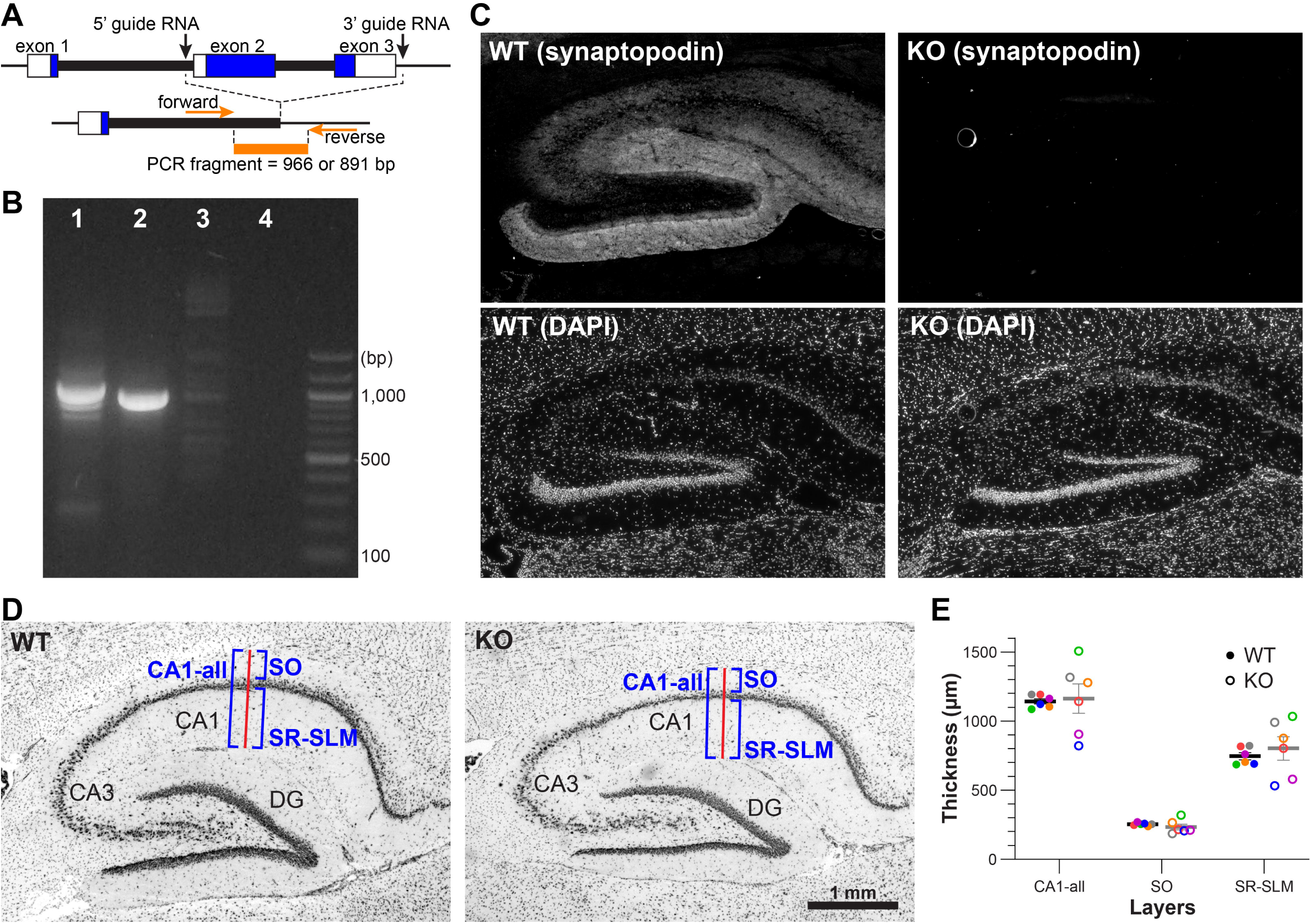
Generation of congenital Synpo KO rat. (A) Schematic of the rat Synpo gene structure. Top: WT gene structure with guide RNA targeting sites (black arrows) flanking exons 2 and 3. Bottom: KO gene structure with locations of forward and reverse PCR primers used to confirm deletion (orange arrows). Blue box = protein-coding exons; white box = untranslated exon regions; thick lines = introns; thin lines = flanking genomic DNA; orange line = expected PCR fragment with successful deletion. (Also see S1 Document.) (B) PCR products from two founder lines show fragments with expected sizes (Lane 1 = Allele A or LE-Synpo*^em1Kmh^*, 966 bp; Lane 2 = Allele B or LE-Synpo*^em2Kmh^*, 891 bp). Lane 3 = WT; PCR band is absent because the conditions do not support amplification of the longer wild-type sequence. Lane 4 = no DNA negative control. The original image of this gel is available in S1 Fig. (C) Synpo protein is absent in the KO hippocampus. (D) Representative brightfield micrographs of the WT (left) and KO (right) hippocampus in parasagittal sections stained with Cresyl Violet. In the area CA1, the thickness of layers (CA1-all, SO, and SR-SLM, delineated by blue brackets) was measured along the red line that originates at the dorsal-most point of the hippocampus and crosses stratum pyramidale perpendicularly, ending at the hippocampal fissure. (E) Quantification shows that CA1 layer thickness was not altered in KO (mean ± SEM; unpaired t-test; n = 6 sections from 3 animals per genotype; colors indicate individual animals). CA1-all: WT = 1142.69 ± 18.62 µm, KO = 1161.73 ± 106.13; p = 0.87. SO: WT = 253.60 ± 4.52, KO = 233.67 ± 20.21; p = 0.22. SLM: WT = 746.10 ± 25.38, KO = 802.88 ± 85.14; p = 0.54.

**Fig 2.**
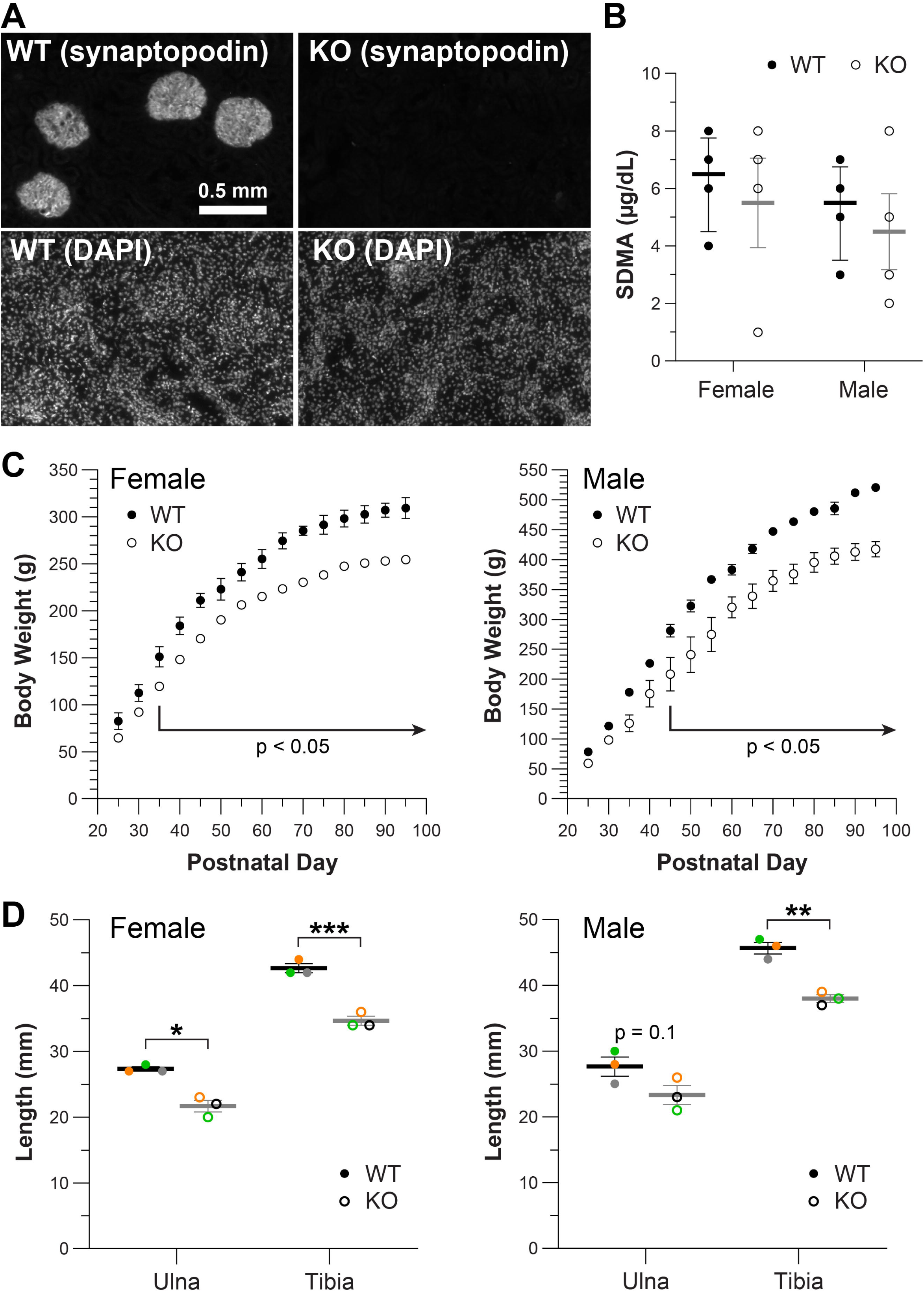
Adult Synpo KO rats show normal excretory renal function, but have a diminished stature compared to WT. (A) Synpo protein is absent in the kidney of KO. (B) Serum SDMA levels are comparable between WT and KO. The median and interquartile range are indicated. p = 0.97 (female) and p = 0.66 (male), Mann-Whitney U test. (C) KO rats had a significantly lower body weight at the age of P35 or older for females and P45 or older for males. N = 3 and mean ± SEM for each data point. Some error bars are smaller than the symbols, and thus too small to be visualized. Two-way analysis of variance (genotype effect for female, F(1, 4) = 167.7, p = 0.0002; male, F(1, 4) = 17.50, p = 0.014 with post hoc Bonferroni’s multiple comparisons). (D) KO rats had significantly shorter ulna and tibia. Mean ± SEM. Female ulna: WT = 27.33 ± 0.33 mm, KO = 21.67 ± 0.88; * p = 0.014. Female tibia: WT = 42.67 ± 0.67, KO = 34.67 ± 0.67; *** p = 0.001. Male ulna: WT = 27.67 ± 1.45 mm, KO = 23.33 ± 1.45; p = 0.1. Male tibia: WT = 45.67 ± 0.88, KO = 38.00 ± 0.58; ** p = 0.003. Unpaired t-test. Colors indicate individual animals.

**Fig 3.**
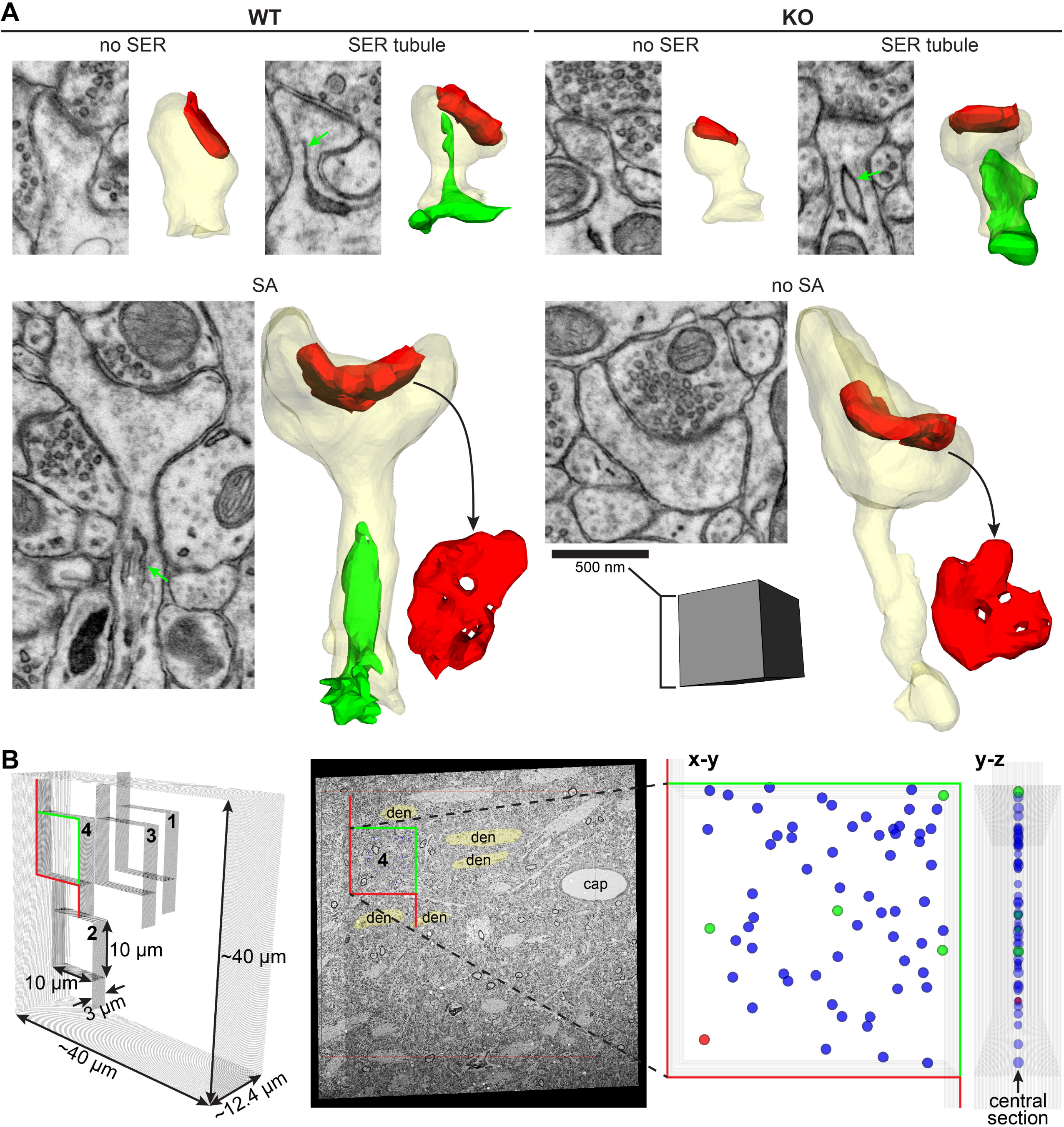
The spine apparatus is absent in Synpo KO rat. (A) EM images and 3D reconstructions of dendritic spines (transparent yellow) and their PSDs (red) in WT and KO, with no SER, SER tubule, or SA (green). KO rats had spines and PSDs as large as those in WT, but no SA were observed. (B) Sample regions of interest (ROI), this example from WT. Left – A schematic view of four ROIs (labeled 1-4; 10 µm × 10 µm × 3 µm each) placed within nonoverlapping locations in one of four EM series (n = 2 per genotype) each encompassing ∼40 µm × ∼40 µm × ∼12 µm. Middle – Each ROI (a sampling frame for ROI #4 is shown) was placed on the central section avoiding large structures such as apical trunk dendrites (den) and capillaries (cap). Right – A schematic view of the ROI #4 in x-y and y-z planes, with colored spheres representing all PSDs that occurred on the central section within the sampling frame. PSDs that cross the green inclusion lines are included, while those crossing the red exclusion lines are excluded from analysis. The identified PSDs were categorized according to the presence of the spine apparatus (red sphere), smooth endoplasmic reticulum (green spheres), or absence of SER or spine apparatus (blue spheres) in the spines. Quantification is summarized in Table 1.

**Fig 4.**
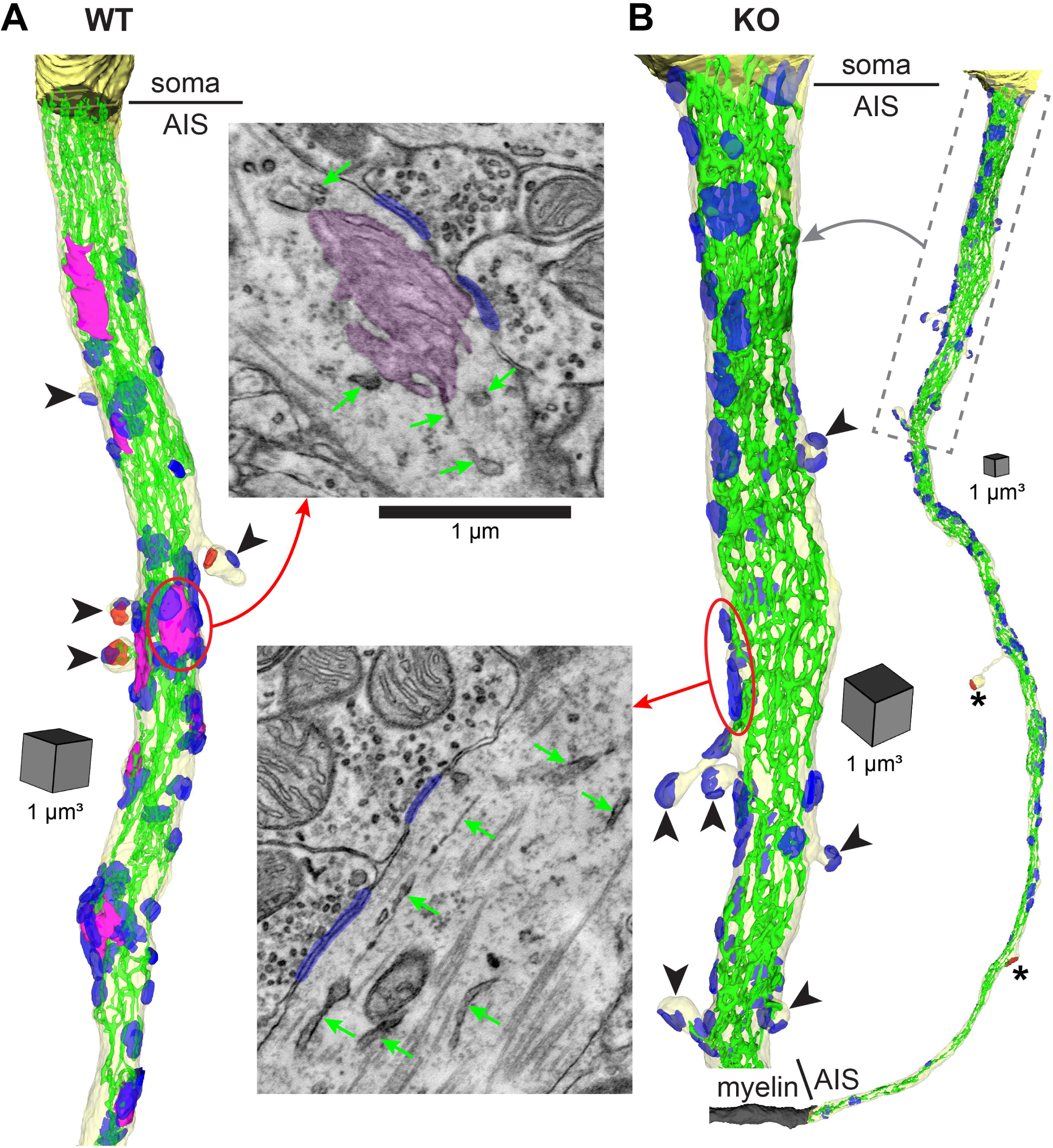
The cisternal organelle is absent from the AIS of *Synpo* KO rats. (A) and (B) EM images and 3D reconstructions of AIS (transparent yellow) and soma (solid yellow; partial reconstructions) with SER (green), cisternal organelles (magenta), symmetric synapses (blue) and asymmetric synapses (red). The AIS from WT (A) had four spines (black arrowheads), three of which are dually innervated by symmetric and asymmetric synapses, while the remaining one was singly innervated by a symmetric synapse. One AIS from KO (B, right) was reconstructed in its entirety from the soma to the beginning of the myelinated axon (gray) and no cisternal organelles were found (see Table 2). Proximal portion of this AIS (gray dotted box) is enlarged to the same scale as WT AIS in A (B, left). This AIS had 15 axonal protrusions, 6 of which are visible and receive inhibitory afferents (indicated by black arrowheads). Asymmetric synapses formed by this AIS onto dendrites are indicated by asterisks.

**Fig 5.**
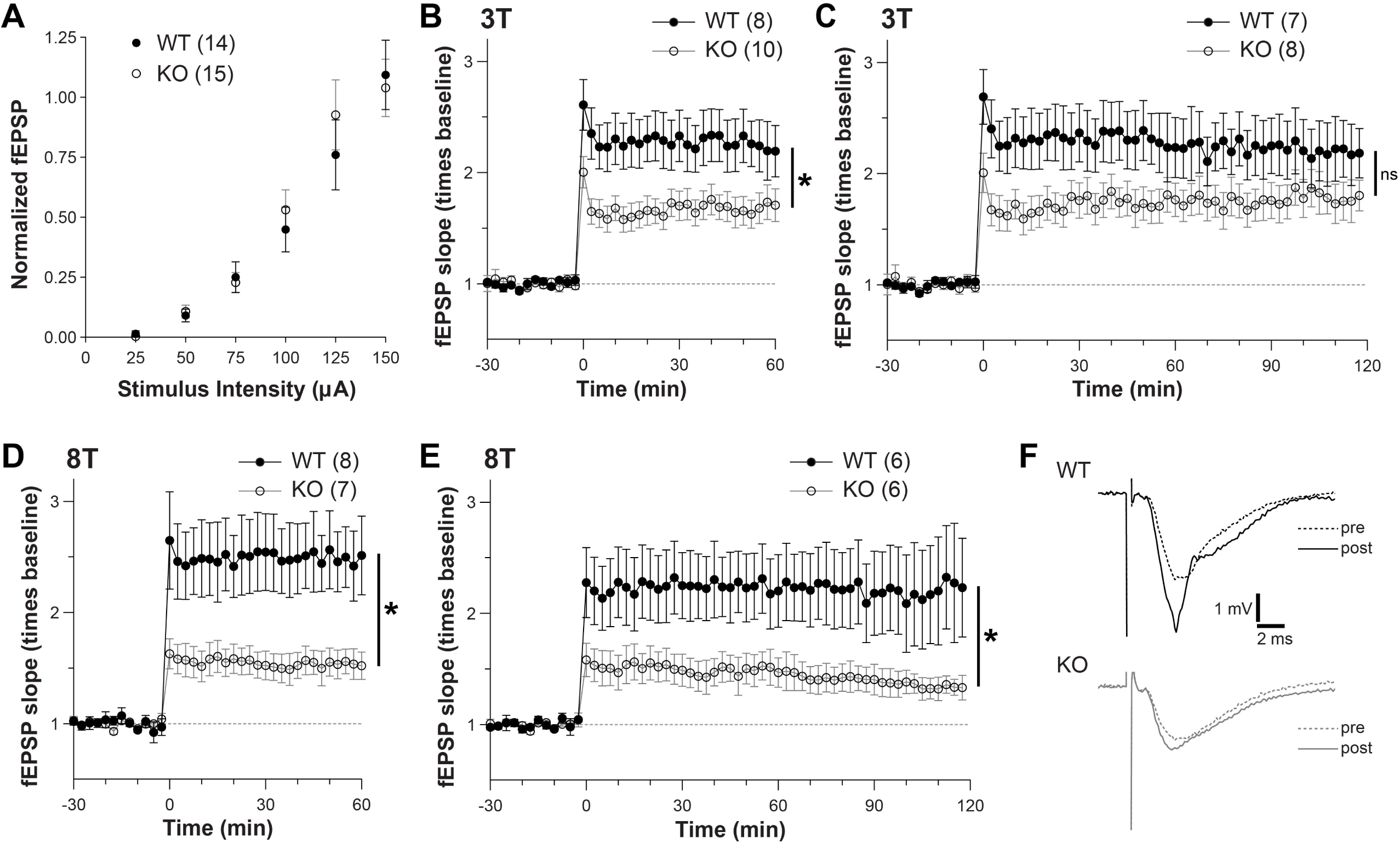
Early LTP is consistently impaired in the *Synpo* KO rats while late-LTP is dependent on the induction paradigm. (A) Overlapping stimulus-response curves for WT and KO. Data were collected prior to baseline recordings during experiments using 3 trains (3T) or 8 trains (8T) of TBS. (B) LTP induced by 3T at time 0 is impaired in Synpo KO at 1 hr after TBS. (WT = 2.29 ± 0.23; KO = 1.68 ± 0.13; *p < 0.05, Mann-Whitney U test.) (C) The difference between WT and KO at 2 hr after 3T is not statistically significant (WT = 2.21 ± 0.27; KO = 1.76 ± 0.1**7**; ns, Mann-Whitney U test). (D) and (E) LTP induced by 8T at time 0 is impaired in Synpo KO at both (D) 1 hr and (E) 2 hr after delivery of 8T. (At 1 hr: WT = 2.49 ± 0.31 and KO = 1.54 ± 0.12. At 2hr: WT = 2.19 ± 0.43 and KO = 1.34 ± 0.09; *p < 0.05, Mann-Whitney U test.) (F) Waveforms from the 8T experiments show example pre- and post-8T responses at 2 hr. Numbers in parentheses above each graph in A-E indicate the number of slices in each experiment. In total, 7 WT and 8 KO rats were used and data from 1-4 slices per animal were included in analyses.

**Table 1.** Quantification of asymmetric axo-spinous synapses and the presence of SER-derived structures in their spines.

| | N | ROI Area ($\mu\text{m}^2$) | PSDs Total Number | PSDs/$\mu\text{m}^2$ (by ROI) | PSDs/$\mu\text{m}^2$ (by animal) | no SER | SER tubule | SA |
| --- | --- | --- | --- | --- | --- | --- | --- | --- |
| WT | 2, 2, 8 | 100 | 473 | $0.59 \pm 0.031$ | 0.55, 0.63 | 419 (89%) | 29 (6%) | 25 (5%) |
| KO | 2, 2, 8 | 100 | 516 | $0.65 \pm 0.029$ | 0.63, 0.66 | 495 (96%) | 21 (4%) | 0 (0%) |

N = number of animals, series, ROIs. PSD density did not differ between WT and KO, mean ± SEM, n = 8 ROIs per genotype, p = 0.22, unpaired t-test.

**Table 2.**
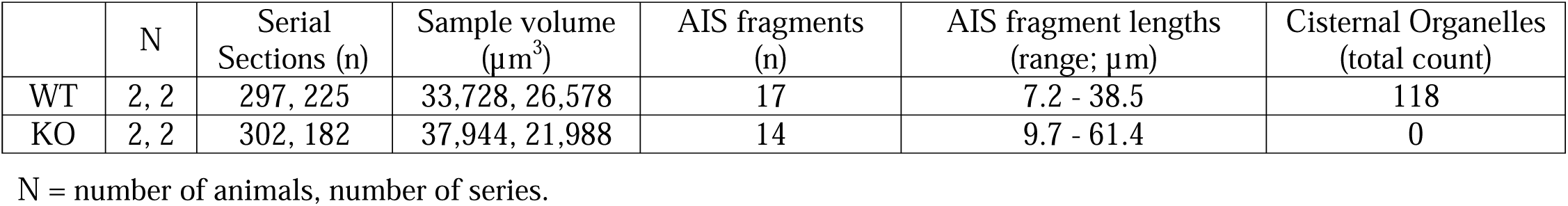
Quantification of AIS and cisternal organelles.

The absence of Synpo protein in the KO was confirmed with immunohistochemistry labeling of the hippocampus (Fig 1C), where Synpo is highly expressed in WT rats. Gross hippocampal morphology was indistinguishable between WT and KO rats. Specifically, there was no statistically significant difference in the thickness of the CA1 layers, including the *stratum oriens* (SO), the combined *strata radiatum* and *lacunosum-moleculare* (SR-SLM), or the entire CA1-all region (Figs 1D and E). Thus, the overall laminar organization of this hippocampal region remains normal in KO rats, a finding consistent with studies in KO mice [1].

Synpo is also expressed in the kidney glomerulus [5,40], where its long and T isoforms modulate Rho signaling and actin bundling in podocytes [18,20], Although their basal urinary protein excretion was reported to be normal [1], *Synpo* KO mice exhibited a delayed recovery from proteinuria during a model of transient nephrotic syndrome [5,21]. The kidney-specific Synpo isoforms are encoded by exons 2 and 3 that were deleted in our rat lines, with their absence in the KO kidney confirmed by immunofluorescence labeling (Fig 2A). We also assessed the levels of symmetric dimethylarginine (SDMA) in blood serum samples collected from WT and KO. SDMA is a byproduct of intranuclear arginine methylation and serves as a biomarker correlating with glomerular filtration rate in multiple mammalian species, including the rat [57–60]. No statistically significant difference was detected in the serum SDMA levels between WT and KO rats (Fig 2B), suggesting that the basal kidney function remains normal in the absence of Synpo. However, over a period of 288 days, 3 KO rats (2 males and 1 female) out of 20 total developed acute renal failure and were euthanized at ages between P105 and P157. During the same period, acute renal failure was not seen in any of the 26 WT or 28 heterozygous littermates P100 or older, or 38 WT and 43 KO that were at P99 or younger. These findings suggest that some KO rats may be more susceptible to conditions of kidney stress later in adulthood.

Unexpectedly, the KO rats of both founder lines appeared consistently smaller in stature than the WT littermates. We tracked their body weight between P21 and P100. For both females and males, KO rats weighed significantly less from P35 onward for females and from P45 onward for males (Fig 2C). We also found that the length of the tibia and ulna in young adult female KO rats (P58-72) were significantly shorter than WT littermates, and the tibia was significantly shorter in KO male rats of the same ages (Fig 2D). Thus, body size differences between genotypes become significant after the first several weeks of postnatal development, a factor that may affect interpretation of behavioral tests that depend on locomotion as a measure of performance.

The spine apparatus and other SER-derived structures were quantified using three-dimensional reconstruction from serial section electron microscopy (3DEM) of axo-spinous synapses in CA1 *s. radiatum* of the WT and KO rat (Fig 3). The spine apparatus was identified as consisting of two or more cisterns of SER with interdigitating dense plates, while an SER tubule in a spine is a membrane-bound structure that is contiguous with the SER network in the parent dendritic shaft (Fig 3A). We adapted the unbiased 3D brick method [61] to determine the relative frequencies of SER tubules and the spine apparatus (Fig 3B). For each of four EM series from *s. radiatum* (2 WT and 2 KO), four non-overlapping sub- volumes (300 µm^3^ each) were selected in the neuropil, avoiding large structures such as cell bodies, capillaries, apical trunk dendrites, myelinated axons, and proximal processes of astrocytes. All postsynaptic densities of axo-spinous synapses that appeared in the middle section of each sub-volume were identified. Then the associated spines were followed to determine the absence or presence of an SER tubule or spine apparatus in the spines (Fig 3B). The density of axo-spinous PSDs was comparable between WT and KO, suggesting that *Synpo* deletion does not disrupt the number of synapses. Most spines in both conditions lack an SER tubule or spine apparatus (Fig 3, Table 1). A single SER tubule occurs in about 6% of WT spines and 4% of KO spines (Fig 3, Table 1). A spine apparatus occurred in 5% of the WT, but none occurred in the KO (Table 1), even when compared qualitatively to spines of similar sizes (Fig 3A).

Using 3DEM, we also sampled the axon initial segments (AIS; n = 17 for WT and 14 for KO) located at the border between CA1 *strata pyramidale* and *oriens.* AISs were identified by visual inspection through serial sections for the presence of fasciculated microtubules and dense undercoating along the intracellular face of the plasma membrane [62,63]. Like the spine apparatus, the cisternal organelles form a complex made of at least two SER cisterns with interdigitating dense plates (Fig 4A, EM). Of the sampled AISs, only one was found in its entirety from its origin at the axon hillock to the first myelin sheath (shown in Fig 4B), while all others appeared in our 3DEM datasets as partial fragments. Their estimated lengths ranged 7-39 µm in the 2 WT series and 9-61 µm in the 2 KO series (Table 2). In these

AISs, a total of 118 cisternal organelles were observed in WT and none in KO (Table 2).. One AIS each from a WT (Fig 4A) and a KO (Fig 4B) was reconstructed, including the plasma membrane, all cisternal organelles, SER, and synapses. The SER cisterns of the WT organelles were continuous with the SER network spanning the entire length of the AIS (Fig 4A). In contrast, the AIS from KO contained no cisternal organelles despite having a robust SER network (Fig 4B).

Regardless of the genotype, putative inhibitory synapses were present along the length of all sampled AISs. These synapses were identified by symmetrically thin presynaptic and postsynaptic specializations and smaller pleomorphic presynaptic vesicles compared to putative excitatory synapses on dendritic spines (compare EM images of synapses in Fig 4 with Fig 3A). Based on a qualitative observation from the reconstructed AIS (Fig 4), the symmetric synapses appeared to cluster around the cisternal organelles in WT, whereas synapses in the KO may be more evenly distributed along the length of the AIS. The reconstructed WT AIS had four spines, of which one was singly innervated by an inhibitory afferent and the remaining three were dually innervated by an excitatory and an inhibitory synapse (Fig 4A). The reconstructed AIS from KO (Fig 4B) had a total of 15 axonal protrusions, of which 10 were spines receiving 1-4 putative inhibitory afferents, four were nonsynaptic, and the remaining one was a presynaptic terminal forming an asymmetric synapse onto an aspiny dendrite (asterisk in Fig 4B). Shaft of this AIS formed another asymmetric synapse onto a separate aspiny dendrite.

Next, we analyzed the impact of *Synpo* deletion on LTP in the rat hippocampal area CA1. Field excitatory postsynaptic potentials (fEPSPs) were recorded in *s. radiatum* of acute hippocampal slices from WT and KO. The stimulus/response curves did not differ significantly between the WT and KO, suggesting basal synaptic transmission was not affected by the *Synpo* deletion (Fig 5A). Stimulus intensities set to ∼50% of maximal yielded comparable baseline fEPSPs across experiments and genotypes (WT = 0.95 ± 0.10 mV/ms; KO = 1.09 ± 0.12; mean ± SEM). A moderate LTP induction protocol, consisting of 3 trains of theta-burst stimulation (TBS, 3T), revealed impaired LTP in slices from KO rats compared to WTs when measured 1 hr after TBS (Fig 5B). However, by 2 hr after 3T the difference in LTP magnitude between WT and KO was no longer statistically significant (Fig 5C). In contrast, a stronger, saturating LTP protocol with 8 trains of TBS produced less LTP at both times in the KO (Figs 5D-F).

## Discussion

We report generation of two rat lines with congenital deletion of both exons 2 and 3 of *Synpo* using a CRISPR- Cas9 approach. Although viable and comparable physically to WT rats at younger ages, adult KO rats exhibit reduced body weight and shortened limbs. Despite normal hippocampal histology, they lack both the spine apparatus and cisternal organelle and display impaired long-term potentiation (LTP). As in KO mice, a subset of KO rats is susceptible to kidney stress in adulthood.

The basal excretory renal function in our rat KO (aged P58-105) is normal based on the SDMA data. In previous studies, dipstick proteinuria was not significantly different between KO and WT mice [5]. One mouse KO model had only exon 2 deleted, and expression of Synpo-T (an isoform encoded by exon 3) was elevated in the kidney of the KO [5].

When these KO mice were challenged with lipopolysaccharides, a model of transient nephrotic syndrome, they recovered, but more slowly than their WT counterparts. The delayed recovery was attributed to a slower re-formation of podocyte actin filaments partially supported by Synpo-T. Another KO mouse model lacking all three Synpo isoforms did not develop kidney abnormalities up to 12 months of age, but they were more susceptible to nephropathy induced by Adriamycin [21]. Since both exons 2 and 3 of the *Synpo* gene are deleted in the KO rat, all three Synpo isoforms are absent, which might contribute to the acute renal failure observed in only 3 KO that were P105 or older. These results suggest that KO rats can be studied during development and in young adulthood without concern for kidney malfunction.

KO rats older than P35 (females) or P45 (males) are smaller in stature than their WT littermates, as measured by body weight. Young adult KO rats also have shorter limb bone length than WT. Additional studies with a larger cohort of animals from wider age range are needed to characterize the KO effect on body development more precisely. It is not known whether the smaller stature of the KO might be secondary to subclinical levels of kidney dysfunction, which indirectly affects calcium reabsorption [64,65], or to a direct role for *Synpo* in musculoskeletal development. An RNA-seq study of 11 organs in Fisher rats (age 2-104 weeks) showed that heart, muscle, and lungs express *Synpo* transcripts at least as much as the brain and kidney, but bone tissue was not included [15]. Another northern blot analysis showed expression of *Synpo* mRNA in the heart, skeletal muscle, and lung of the rat [14]. Previous reports from KO mice mentioned that their appearance was indistinguishable from WT but did not measure body weight and bone dimensions even though they might be affected [1]. Furthermore, the small size of mice makes it harder to detect and measure subtle but important differences in bone length and body mass. Mice also express *Synpo* transcripts in the heart and lungs at 8 weeks, and limbs at embryonic day 14.5 [16]. In addition, a recent study confirmed expression of both Synpo transcripts and proteins in the striated and smooth muscles in mice [17]. Thus, the potential involvement of *Synpo* in limb maturation could confound behavioral analyses, such as the reduced horizontal locomotor activity observed for KO mice in the open-field test [1]. Body weight may also affect performance in some motor tasks such as rotarod and rope climbing [66,67]. A more refined approach to analyzing microstructure of behavior using the Behavioral Pattern Monitor [68] may also help characterize in detail the potential KO effects on motor performance in both rats and mice. Being able to assess when during development the gross body effects occur in the KO is thus an advantage for the rat KO model system.

SER is a continuous internal membrane that extends from the cell body into axons, dendrites, and some spines [42,69–72]. SER regulates calcium dynamics and homeostasis, and the synthesis and trafficking of lipids and proteins [73,74]. SER is a limited resource that makes brief visits to active spines [75], which may explain why SER occurs in less than 15% of hippocampal CA1 spines at any one time [33,42,43]. We found a comparable proportion of dendritic spines contains an SER tubule in both KO and WT rats (at P80), suggesting the capacity for SER to visit spines is not lost in the *Synpo* KO.

The spine apparatus is a specialized structure derived from SER and dense plates containing Synpo [3,76]. In addition to its SER-related functions, the spine apparatus is often associated with ribosomes and polyribosomes, indicating a role in local protein synthesis [26,28]. Secretory pathway proteins found in the Golgi apparatus are also present in the spine apparatus, suggesting involvement in local post-translational modifications critical for maturation and correct trafficking of proteins [27]. In the adult hippocampus, the largest synapses occur on dendritic spines that contain SER, especially in the form of the spine apparatus. By 2 hours after the induction of LTP, the predominant type of SER contained in spines shifts from a single tubule under control conditions to a fully elaborated spine apparatus [33]. LTP induces dramatic PSD enlargement in spines with SER, while spines without SER undergo a comparatively smaller increase in PSD size. In addition, dendritic spines tend to cluster in the vicinity of large spines that contain SER. Intracellular trafficking slows where the dendritic shaft SER branches or expands, allowing ER exit sites to locally deliver resources that support clustered synapses [77]. High density spine clusters are longer and have more SER branches than low density clusters. Following LTP, the density of spines in a cluster that has an SER-containing spine remains high and total synapse area is elevated as these synapses enlarge. In contrast, spine density in a cluster without SER-containing spines remains low or is further reduced following LTP. Furthermore, high-density clusters display fewer SER branches in the dendritic shaft post-LTP, possibly reflecting migration of SER into spines to support PSD growth and elaboration of the spine apparatus [33]. The spine apparatus is eliminated in the *Synpo* KO, yet qualitative observations suggest that spines of comparable sizes could be found in the KO and WT of our rat and in the mouse [1]. Future work will be needed to determine whether spines and synapses reach maximal size in the KO and whether clustering of small spines around enlarged spines can be enhanced during plasticity.

Axo-axonic GABAergic interneurons, also known as Chandelier cells, selectively target the AIS of excitatory pyramidal neurons in hippocampal area CA1 and other excitatory pyramidal neurons throughout the brain. This network is important in many normal brain functions such as slow wave sleep and circuit excitability [78]. Dysfunction of the axo-axonic network has been linked to schizophrenia, epilepsy, and autism spectrum disorder [79]. Synpo is required for the formation of cisternal organelles in the AIS of mice [2]. Hence, the absence of the cisternal organelle in the AIS of the CA1 pyramidal cells in our adult KO rat is expected. A previous report from the mouse neocortical pyramidal neurons showed that inhibitory synapses cluster around the cisternal organelles [80]. Our qualitative observation based on a reconstructed AIS suggests that this may also be the case in the rat hippocampal area CA1. Inhibitory synapses were still present along the AIS in our KO rats, but they appear to be distributed more uniformly. These initial observations motivate future quantitative studies, including cluster analyses, of how Synpo regulates the spatial relationship between the cisternal organelles and inhibitory synapses along AISs. Although LTP in CA1 is usually associated with glutamatergic synapses and plasticity in other interneuron types (including those targeting dendrites or soma), the spatial arrangement of axo-axonic synapses at the AIS could indirectly influence plasticity by modulating pyramidal neuron firing patterns [81].

Multiple factors could contribute to the differential impact of *Synpo* deletion on LTP in KO mice versus KO rats.

In mice, the greater neuronal density compared to rats can affect synaptic connectivity, network activity, neuronal excitability, as well as the expression of synaptic plasticity [82–85]. We evaluated LTP induction and maintenance in adult KO rats using 3T and 8T TBS protocols consistent with one of the original KO mouse studies and our previous rat studies [1,38,86]. Both protocols result in reduced LTP in the *Synpo* KO rat at 1 hr after induction. LTP remains impaired at 2 hr after induction by 8T but not by 3T in the KO rats, contrasting with the KO mice that remained impaired at 2 hr after 3T [1]. LTP induction protocols varied greatly across KO mouse studies including tetanic stimulation at 100 Hz [1,46,48] versus TBS for 1T, 3T, or 5T [1,45]. These distinct stimulation patterns could activate different molecular pathways [87] and recruit varying numbers of presynaptic axons. Age also plays an important role in LTP ranging from P15 to 6 months for the various studies of LTP in KO mice. Zhang et al. showed, for example, that LTP was impaired in juvenile but not adult KO mice [45]. The dramatic difference between the normal developmental onset and maturation of

LTP across ages P18-P37 in mice [51] contrasts with the discrete onset age at P12 in the rat [53], which likely contributes to the differential effects between mice and rats of *Synpo* KO on the expression of LTP. The *Synpo* KO rat could serve as a valuable tool to assess whether the developmental upregulation of Synpo [40,41] and formation of the spine apparatus [42,43] are necessary for the maturation of synapses, spine clustering, and LTP [33,39].

The spine apparatus plays a critical role in human health and disease, as demonstrated by its aberrant morphology in neurodegenerative and peritumoral brain tissue [88]. Downregulation of Synpo has also been linked to Alzheimer’s disease, inflammation, stress, and embryonic hypoxia [9,89–93]. Emerging evidence also supports the use of Synaptopodin as a biomarker for other neural diseases, including frontotemporal dementia, multiple sclerosis, and acute brain injury [8,10,12].

The effect of *Synpo* deletion on bone development and body growth could be mitigated by a tissue- and time- specific deletion strategy, using, for example, the Cre-lox recombination system [94,95]. Such an approach would allow the animals to develop normally into adulthood before the gene of interest is excised in an inducible manner. However, an inducible KO model is not practical to assess the role of Synpo during development because the Synpo protein has a half- life of about 10 days and is expressed soon after birth by P5, which is well before the spine apparatus forms [41,42,96].

Our congenital KO rat is thus a valuable resource for investigating the function of Synpo during the first 40 postnatal days of life when animals show little or no confounding kidney, bone, or weight gain defects.

## Materials and Methods

This study was carried out in accordance with the US National Research Council’s Guide for the Care and Use of Laboratory Animals, the US Public Health Service’s Policy on Humane Care and Use of Laboratory Animals, and the Guide for the Care and Use of Laboratory Animals. All animal procedures were approved by the Institutional Animal Care and Use Committee at The University of Texas at Austin. All rats were housed under 12-hr light/dark cycles with water and food available *ad libitum* in an AAALAC-accredited facility managed by the UT-Austin Animal Resource Center. All efforts were made to minimize suffering.

### Congenital *synaptopodin* knockout lines

The *Synpo* KO lines were generated at Mouse Genetic Engineering Facility at UT-Austin (RRID:SCR_021927) using an approach based on the CRISPR-Cas9 system. Synthetic guide RNAs were designed to target 5’ and 3’ regions flanking the protein-coding regions of the *Synpo* gene, exons 2 and 3, using CRISPOR (version 4.7; https://crispor.org; RRID:SCR_015935; [97]) (Fig 1A and S1 Document). These guide RNAs were predicted by CRISPOR to be highly specific, and there are no potential off-target sequences in known exons or on chromosome 18, where *Synpo* gene is located. Using well-designed guide RNAs should make off-target mutations rare and indistinguishable in frequency from naturally occurring mutations [98]. Should there be such an off-target mutation on another chromosome, it is likely to be lost during additional breeding since that chromosome is not being selected for. The efficacy of the guide RNAs was tested in cultured rat embryos, and the best ones were then used *in vivo*. The guide RNA was delivered by microinjection of a ribonucleoprotein containing a high-fidelity Cas9 enzyme (HiFi Cas9 nuclease, IDT) for short half-life to reduce the editing window. Injection of the 5’ and 3’ guide RNAs together into rat embryos resulted in two founder lines with congenital *Synpo* deletion (Fig 1B, lanes 1 and 2). These lines were bred and maintained separately because their alleles differ in the genomic location of the breakpoints (see S1 Document). Genotypes of the animals were confirmed before weaning (P10-17) and again at the time of euthanasia for experiments (genotyping performed by Transnetyx, Cordova, TN). The lines were generated and maintained in the Long-Evans background (Charles River strain 006; Crl:LE; RRID:RGD_2308852), and a new WT rat was introduced about every 5 generations to prevent genetic drift. The KO lines are designated as LE-Synpo*^em1Kmh^*(RRID:RRRC_00964) and LE-Synpo*^em2Kmh^* (RRID:RRRC_01025) and have been registered with the Rat Genome Database (https://rgd.mcw.edu) as RGD ID 155782907 and RGD ID 616335891, respectively.

### Vaginal smear cytology

All adult female rats were used during diestrus when circulating estradiol is low and relatively stable. The vaginal smear cytology was measured on the day of the experiment and several days prior. The estrous cycle of each rat was monitored by performing daily vaginal lavage using 20 µl sterile saline and recording cell morphology on glass slides under an upright microscope at 10× magnification. Diestrus is characterized by the highest proportion of leukocytes [99–101].

### Brain and kidney tissue for light microscopy

For immunofluorescence labeling and histology staining, the rats (n = 2 females and 1 male per genotype; P80; LE-Synpo*^em2Kmh^*) were rapidly decapitated under heavy isoflurane anesthesia. The brain was removed from the cranium, rinsed with phosphate-buffered saline (PBS), and bisected along the midline. Both hemispheres were placed on a metal spatula with the cut surface down and frozen in 2-methyl butane cooled with dry ice. The kidney was also dissected out and frozen as a whole in the same manner. The frozen tissue was stored at -80°C until being cut into sections at 20 µm thickness using a cryostat. The brain sections were cut in the parasagittal plane. The sections were mounted on glass microscope slides and were stored at -80°C until they were used for immunolabeling or histology staining.

### Immunofluorescence labeling

The slide-mounted sections were thawed and fixed in 4% formaldehyde in 0.1 M phosphate buffer (PB; pH = 7.4) for 20 min at RT. After PBS washes, the sections were incubated with blocking solution (Animal-Free Blocker® and Diluent; Vector Laboratories, Newark, CA; catalog# SP-5035) containing 0.3% Triton X-100 for 1 hr. The sections were then incubated with rabbit anti-synaptopodin (1:500; Synaptic Systems, Göttingen, Germany; catalog# 163002; RRID:AB_887825) in PBS with the Vector blocking reagent and 0.3% Triton X-100 for overnight at RT. Following PBS washes, the sections were incubated with goat anti-rabbit IgG conjugated with Alexa Fluor 488 (1:100; Thermo Fisher Scientific, Waltham, MA; catalog# A-11034; RRID:AB_2576217) in the same diluent for 1 hr at RT. The sections were washed with PBS and stained with 4′,6-diamidino-2-phenylindole (DAPI) for 5 min. Following washes with PBS and saline (0.9% NaCl, aq), coverslips were applied with Aqua-Poly/Mount anti-fade mountant (Polysciences, Warrington, PA; catalog# 18606). In control experiments, the primary antibody was omitted to confirm the lack of non-specific binding of the secondary antibody, and both the primary and secondary antibodies were omitted to assess background autofluorescence. The primary antibody was raised against a recombinant peptide corresponding to amino acids 331–452 of the mouse Synpo that is conserved in the rat. This antibody recognizes all isoforms of Synpo, and its specificity has been validated by Western blot with brain tissue from the *Synpo* KO mouse by the manufacturer.

Epifluorescence images were acquired with a 2.5× objective on a Zeiss Axio Observer 5 microscope equipped with a Zeiss Axiocam 705 CMOS camera controlled by the ZEN software (version 3.2; RRID:SCR_013672). The Synpo signal was visualized with a filter set for excitation at 450-490 nm and emission at 500-550 nm. The DAPI signal was visualized with a filter set for excitation at 365 nm and emission at 420-470 nm. For both channels, the exposure time was determined by imaging the hippocampus in a Synpo-labeled WT section and applied to all sections from both genotypes.

### Histology

The slide-mounted sections were thawed and fixed in 4% formaldehyde as described above. After being washed in purified water, the sections were stained with Cresyl Violet (FD NeuroTechnologies, Columbia, MD; catalog# PS102- 01). Following dehydration in ascending grade of ethanol and clearing in xylene, coverslips were applied with DPX mountant. Brightfield images encompassing the hippocampus and the overlaying neocortex were acquired with a 2.5× objective on the same Zeiss Axio Observer 5 microscope using Köhler illumination.

The thickness of layers in the dorsal hippocampal area CA1 was measured from images of parasagittal sections (2 per animal) that best corresponded to 1.4 mm lateral according to a rat brain atlas [102]. FIJI (https://fiji.sc/; RRID:SCR_002285; [103]) was used for the measurement along the line that intersects the most dorsal point of the hippocampus and perpendicular to *stratum pyramidale*, from the border between alveus and *stratum oriens* to the hippocampal fissure. Four individual experimenters independently performed measurements masked as to the genotype and the measured thickness was averaged per layer per section.

### 3DEM

For electron microscopy, the rats (n = 2 females per genotype; P80; both LE-Synpo*^em1Kmh^* and LE-Synpo*^em2Kmh^*) were perfusion-fixed under deep isoflurane anesthesia and tracheal supply of oxygen [104]. The perfusion procedure involved a brief (up to ∼20 s) wash with oxygenated Krebs-Ringer Carbicarb buffer (concentration in mM: 2.0 CaCl_2_, 11.0 D-glucose, 4.7 KCl, 4.0 MgSO_4_, 118 NaCl, 12.5 Na_2_CO_3_, 12.5 NaHCO_3_; pH 7.4; osmolality 300-330 mmol/kg), followed by fixative containing 2.0% formaldehyde, 2.5% glutaraldehyde, 2 mM CaCl_2_, and 4 mM MgSO_4_ in 0.1 M cacodylate buffer (pH 7.4) for ∼1 hr, during which ∼1.9 L of fixative was used per animal (both aldehydes from Ladd Research, Essex Junction, VT). The brains were removed from the skull about 1 hr after the end of perfusion, and stored in the same fixative overnight at RT.

The perfusion-fixed brain tissue was cut into parasagittal sections (250 µm thickness) with a Leica VT1000S vibrating blade microtome. After washes in 0.15 M sodium cacodylate buffer (pH = 7.4), the tissue containing the dorsal hippocampus was stained *en bloc* with reduced osmium (2% osmium tetroxide and 3% potassium ferrocyanide in 0.15 M cacodylate buffer) for 45 min followed by washes in the buffer. The tissue was then treated with 320 mM pyrogallol (aq) for 1 hr. After washes in the buffer, the tissue was stained with 2% osmium tetroxide for 45 min, followed by washes in the buffer and purified water. The tissue was then dehydrated in an ascending series of acetonitrile (25%, 50%, 75%, and twice in 100%). The 50% and 75% acetonitrile solutions also contained 2% uranyl acetate for *en bloc* staining for 20 min each. The tissue was infiltrated and embedded into low-viscosity epoxy resin composed of EMBed-812, nonenyl succinic anhydride, nadic methyl anhydride, and benzyldimethylamine at 60°C for 48-60 hr (all resin components from Electron Microscopy Sciences [EMS], Hatfield, PA; [105]). The resin embedded tissue was trimmed to the surface of ∼80 µm × 800 µm, exposing the area CA1 of the dorsal hippocampus encompassing the *s. oriens* and *s. radiatum*. Then, 182-302 serial ultrathin sections (nominal thickness of 45 nm) were obtained using a DiATOME 35° diamond knife on a Leica UC7 ultramicrotome. The sections were collected using Synaptek Be-Cu slot grids (EMS or Ted Pella, Redding, CA) that were coated with polyetherimide.

For analysis of the spine apparatus, the serial sections were imaged from the middle of *s. radiatum* (image field centered at ∼125 µm from the apical face of *s. pyramidale*), masked as to condition, with a transmission-mode scanning EM (tSEM; Zeiss SUPRA 40 field-emission SEM with a retractable multimode transmitted electron detector and ATLAS package; [106]) For the cisternal organelle, the image field centered around the border between *strata oriens* and *pyramidale*. The scan beam at the accelerating voltage of 28 kV was set for a dwell time of 1.3 µs. Each section was imaged from a single field encompassing 49.155 µm × 49.155 µm (24,576 pixels × 24,576 pixels at 2 nm/pixel). The serial tSEM images were aligned using signal whitening Fourier transform [107] deployed at 3dem.org. EM series with the aligned images were assigned a 5-letter code to mask the identity and genotype of the animal and imported into PyReconstruct (https://github.com/synapseweb/pyreconstruct; RRID:SCR_027562; [108]). The pixel size was calibrated using a grating replica (EMS, catalog# 80051), and the average section thickness was calculated with the cylindrical diameter method [109].

### Electrophysiology

Male rats (P80-172; n = 7 for WT and n = 8 for KO) from the line LE-Synpo*^em2Kmh^* were used to prepare acute hippocampal slices. None of the KO rats used in these experiments showed any signs of renal failure. The animals were deeply anesthetized with isoflurane and decapitated with guillotine. The brain was removed from the cranium, and the left hippocampus was dissected out and rinsed with room temperature artificial cerebrospinal fluid (aCSF) containing (in mM) 117 NaCl, 5.3 KCl, 26 NaHCO_3_, 1 NaH2PO_4_, 2.5 CaCl_2_, 1.3 MgSO_4_, and 10 D-glucose, pH 7.4, and bubbled with 95% O_2_-5% CO_2_. Slices (400 μm thick) from the dorsal hippocampus were cut at 70° transverse to the long axis on a Stoelting tissue chopper and transferred in oxygenated aCSF to the 4 chambers in the Synchroslice system (Lohmann Research Equipment, Castrop-Rauxel, Germany). The dissection and slice preparation took <6 min, which is crucial timing for enduring LTP that lasts >3 hr [110]. Hippocampal slices were placed on a net at the liquid-gas interface between 32-33°C aCSF and humidified 95% O_2_-5% CO_2_ atmosphere. After 3 h of incubation, the stimulating and recording electrodes were positioned ∼400 μm apart in the middle of hippocampal CA1 *s. radiatum* with the stimulating electrode on the CA3 side. Stimuli consisted of 200 μs biphasic current pulses, lasting 100 μs each for positive and negative components of the stimulus. Test pulses (50-250 μA) were given at 1 pulse per 2.5 min unless stated otherwise, and field excitatory postsynaptic potentials (fEPSPs) were recorded. The initial fEPSP slope was ∼50% of the maximal fEPSP slope based on the input/output curve for each slice. LTP was induced by three (3T) or eight trains (8T) of TBS with 30 s intervals. Each train of TBS contained 10 bursts at 5 Hz, and each burst contained 4 pulses at 100 Hz.

The delivery of presynaptic stimulation and the acquisition and analysis of fEPSPs were performed with SynchroBrain software (Lohmann Research Equipment). The initial maximum slope was measured over a 0.3-0.5 ms period with position on the waveform held constant in each slice. To calculate the magnitude of LTP, the average of fEPSP slopes during the last 30 min of baseline recordings before the delivery of the TBS was computed and then compared to the average values during the last 20 min of each 1 h or 2 h session following delivery of TBS. Then values across slices (means ± SEM) were presented as times baseline, where 1 indicated no change in the fEPSP slope relative to the pre-TBS baseline. Up to four slices could be recorded simultaneously from a single animal. Typically, two LTP induction protocols (3T and 8T) were performed in the same animal. Each slice was treated as an independent data point for the LTP analysis, and values were averaged across slices within each protocol group (see additional data deposited at the Texas Data Repository, indicated under “Data and code availability”). For each recorded slice, LTP was calculated as the mean of 9 normalized fEPSP values obtained during the last 20 min of recording (40–60 min frame for 1 h LTP and 97.5–117.5 min frame for 2 h LTP). For each genotype-protocol combination, the individual means were averaged across slices.

### Blood sample collection

Blood samples (100 µl per animal) were collected from rats (P58-105, n = 16, 4 per genotype per sex; LE- Synpo*^em2Kmh^*) immediately after euthanasia and were centrifuged at 2500 rpm for 15 min. The resulting serum sample was shipped overnight on ice to be analyzed for symmetric dimethylarginine (SDMA) levels by IDEXX (Westbrook, ME).

### Anatomical measurements

The body weight was measured three times a week from P25 to P100 (n = 6 each for male WT and KO; n = 6 for female WT and n = 3 for female KO; both LE-Synpo*^em1Kmh^* and LE-Synpo*^em2Kmh^*). Because not all animals were weighed on the same days, data was averaged into 4-day bins, with each bin containing measurements for 3 animals. Not all bins contained the same 3 rats.

For measuring bone length, the forelimb and hindlimb were removed from euthanized rats (P58-72, n = 12, 3 per sex per genotype; LE-Synpo*^em2Kmh^*). The skin and large muscles were removed with surgical tools, and the remaining soft tissue was dissolved by overnight incubation with agitation at 50°C in a 5% aqueous solution of laundry detergent containing enzymes and oxygen bleach (Pro-Enzyme Laundry Detergent, Absolute Best Cleaning Products LLC, Brookings, SD). The bones were then rinsed in water, and the length was measured from the ulna and tibia using calipers.

### Statistics

GraphPad Prism (https://www.graphpad.com/; RRID:SCR_002798) was used for statistical tests described in each figure. All data are reported as mean ± SEM unless otherwise noted.

## Resource availability

### Lead contact

Requests for further information and resources should be directed to and will be fulfilled by the lead contact, Masaaki Kuwajima.

### Materials availability

Rat lines generated in this study have been deposited to the Rat Resource and Research Center (Columbia, MO; https://www.rrrc.us), under the designations LE-Synpo*^em1Kmh^*(RRID:RRRC_00964) and LE-Synpo*^em2Kmh^* (RRID:RRRC_01025).

### Data and code availability

- Supporting Information (S1 Document and S1 Fig), 3DEM dataset (original EM images and PyReconstruct files) and spreadsheet files containing data on the hippocampal anatomy (LM and EM), SDMA levels, body weight, bone length, and electrophysiology have been deposited at the Texas Data Repository (https://doi.org/10.18738/T8/HDOTFV) and 3DEM.org (https://3dem.org/public-data/tapis/public/cloud.data/corral-repl/projects/NeuroNex-3DEM/Public/2025_Kuwajima_Synpo-KO-rats/), and they are publicly available as of the date of publication.
- This paper does not report original code.
- Any additional information required to reanalyze the data reported in this paper is available from the lead contact upon request.

## Acknowledgments

This work was supported by the Mouse Genetic Engineering Facility (RRID:SCR 021927; Dr. William Shawlot), a core facility within the Center for Biomedical Research Support at the University of Texas at Austin. The authors thank Dr. Katy Pilarzyk Alvarado for conducting preliminary work on this study, Patrick Parker, Myles Joyce, Isha Jha, and Everett Owens for data collection, Anna-Maria Escherich for administrative and animal colony support, and Dr. Michael Drew for the use of a cryostat.

## Author Contributions

Conceptualization: MK, LMK, KMH Funding acquisition: KMH

Methodology: MK, OIO, LMK, SS, KMH

Investigation: MK, OIO, AA, WY, SS, AX, AL, EP

Data curation: MK, OIO, WY, KMH

Formal analysis: MK, OIO, AA, WY, SS, AX, AL, EP, KMH

Visualization: MK, OIO, LMK, AA, WY, SS, AX, AL, EP

Writing – original draft: MK, OIO, SS

Writing – review and editing: MK, OIO, LMK, WY, KMH

Supervision and Project administration: MK, KMH

## Declaration of Generative AI and AI-assisted Technologies

The authors did not use generative AI and AI-assisted technologies during the preparation of this work.

## Supporting Information

**S1 Document. Information on CRISPR Reagents and Genotyping. S1 Figure. Original gel image used in Figure 1B**.

## Additional Resources

The Harris lab wiki describing details of 3DEM procedures (perfusion-fixation, tissue processing, and imaging): https://cloud.wikis.utexas.edu/wiki/spaces/khlab/

## References

1. Deller T, Korte M, Chabanis S, Drakew A, Schwegler H, Stefani GG, et al. Synaptopodin-deficient mice lack a spine apparatus and show deficits in synaptic plasticity. Proc Natl Acad Sci U S A. 2003;100: 10494–10499. doi:10.1073/pnas.1832384100

2. Bas Orth C, Schultz C, Müller CM, Frotscher M, Deller T. Loss of the cisternal organelle in the axon initial segment of cortical neurons in synaptopodin-deficient mice. J Comp Neurol. 2007;504: 441–449. doi:10.1002/cne.21445

3. Falahati H, Wu Y, Fang M, De Camilli P. Ectopic reconstitution of a spine-apparatus-like structure provides insight into mechanisms underlying its formation. Curr Biol. 2024 [cited 10 Dec 2024]. doi:10.1016/j.cub.2024.11.010

4. Korkotian E, Segal M. Synaptopodin regulates release of calcium from stores in dendritic spines of cultured hippocampal neurons. J Physiol. 2011;589: 5987–5995. doi:10.1113/jphysiol.2011.217315

5. Asanuma K, Kim K, Oh J, Giardino L, Chabanis S, Faul C, et al. Synaptopodin regulates the actin-bundling activity of α-actinin in an isoform-specific manner. J Clin Invest. 2005;115: 1188–1198. doi:doi:10.1172/JCI23371

6. Hirakawa M, Tsuruya K, Yotsueda H, Tokumoto M, Ikeda H, Katafuchi R, et al. Expression of synaptopodin and GLEPP1 as markers of steroid responsiveness in primary focal segmental glomerulosclerosis. Life Sci. 2006;79: 757–763. doi:10.1016/j.lfs.2006.02.031

7. Kwon SK, Kim SJ, Kim H-Y. Urine synaptopodin excretion is an important marker of glomerular disease progression. Korean J Intern Med. 2016;31: 938–943. doi:10.3904/kjim.2015.226

8. Goetzl EJ, Kapogiannis D, Schwartz JB, Lobach IV, Goetzl L, Abner EL, et al. Decreased synaptic proteins in neuronal exosomes of frontotemporal dementia and Alzheimer’s disease. FASEB J. 2016;30: 4141–4148. doi:10.1096/fj.201600816R

9. Datta A, Chai YL, Tan JM, Lee JH, Francis PT, Chen CP, et al. An iTRAQ-based proteomic analysis reveals dysregulation of neocortical synaptopodin in Lewy body dementias. Mol Brain. 2017;10: 36. doi:10.1186/s13041-017-0316-9

10. Goetzl L, Merabova N, Darbinian N, Martirosyan D, Poletto E, Fugarolas K, et al. Diagnostic Potential of Neural Exosome Cargo as Biomarkers for Acute Brain Injury. Ann Clin Transl Neurol. 2017;5: 4–10. doi:10.1002/acn3.499

11. Winston CN, Goetzl EJ, Baker LD, Vitiello MV, Rissman RA. Growth Hormone-Releasing Hormone Modulation of Neuronal Exosome Biomarkers in Mild Cognitive Impairment. J Alzheimers Dis JAD. 2018;66: 971–981. doi:10.3233/JAD-180302

12. Bhargava P, Nogueras-Ortiz C, Kim S, Delgado-Peraza F, Calabresi PA, Kapogiannis D. Synaptic and complement markers in extracellular vesicles in multiple sclerosis. Mult Scler J. 2021;27: 509–518. doi:10.1177/1352458520924590

13. Ladakis DC, Vreones M, Blommer J, Harrison KL, Smith MD, Vasileiou ES, et al. Synaptic Protein Loss in Extracellular Vesicles Reflects Brain and Retinal Atrophy in People With Multiple Sclerosis. Neurol Neuroimmunol Neuroinflammation. 2024;11: e200257. doi:10.1212/NXI.0000000000200257

14. Yamazaki M, Matsuo R, Fukazawa Y, Ozawa F, Inokuchi K. Regulated expression of an actin-associated protein, synaptopodin, during long-term potentiation. J Neurochem. 2001;79: 192–199. doi:10.1046/j.1471-4159.2001.00552.x

15. Yu Y, Fuscoe JC, Zhao C, Guo C, Jia M, Qing T, et al. A rat RNA-Seq transcriptomic BodyMap across 11 organs and 4 developmental stages. Nat Commun. 2014;5: 3230. doi:10.1038/ncomms4230

16. Yue F, Cheng Y, Breschi A, Vierstra J, Wu W, Ryba T, et al. A comparative encyclopedia of DNA elements in the mouse genome. Nature. 2014;515: 355–364. doi:10.1038/nature13992

17. Lohanadan K, Assent M, Linnemann A, Qu C, Schänzer A, Heukamp L, et al. Characterization of synaptopodin in striated and smooth muscles: isoform spectrum, expression patterns, localization and protein interactions. Exp Cell Res. 2026; 114990. doi:10.1016/j.yexcr.2026.114990

18. Asanuma K, Yanagida-Asanuma E, Faul C, Tomino Y, Kim K, Mundel P. Synaptopodin orchestrates actin organization and cell motility via regulation of RhoA signalling. Nat Cell Biol. 2006;8: 485–491. doi:10.1038/ncb1400

19. Faul C, Donnelly M, Merscher-Gomez S, Chang YH, Franz S, Delfgaauw J, et al. The actin cytoskeleton of kidney podocytes is a direct target of the antiproteinuric effect of cyclosporine A. Nat Med. 2008;14: 931–938. doi:10.1038/nm.1857

20. Buvall L, Wallentin H, Sieber J, Andreeva S, Choi HY, Mundel P, et al. Synaptopodin Is a Coincidence Detector of Tyrosine versus Serine/Threonine Phosphorylation for the Modulation of Rho Protein Crosstalk in Podocytes. J Am Soc Nephrol. 2017;28: 837–851. doi:10.1681/ASN.2016040414

21. Ning L, Suleiman HY, Miner JH. Synaptopodin Is Dispensable for Normal Podocyte Homeostasis but Is Protective in the Context of Acute Podocyte Injury. J Am Soc Nephrol. 2020;31: 2815–2832. doi:10.1681/ASN.2020050572

22. Sánchez-Ponce D, Blázquez-Llorca L, DeFelipe J, Garrido JJ, Muñoz A. Colocalization of α-actinin and Synaptopodin in the Pyramidal Cell Axon Initial Segment. Cereb Cortex. 2012;22: 1648–1661. doi:10.1093/cercor/bhr251

23. Jungenitz T, Bird A, Engelhardt M, Jedlicka P, Schwarzacher SW, Deller T. Structural plasticity of the axon initial segment in rat hippocampal granule cells following high frequency stimulation and LTP induction. Front Neuroanat. 2023;17. doi:10.3389/fnana.2023.1125623

24. Vlachos A, Maggio N, Segal M. Lack of correlation between synaptopodin expression and the ability to induce LTP in the rat dorsal and ventral hippocampus. Hippocampus. 2008;18: 1–4. doi:10.1002/hipo.20373

25. Holbro N, Grunditz Å, Oertner TG. Differential distribution of endoplasmic reticulum controls metabotropic signaling and plasticity at hippocampal synapses. Proc Natl Acad Sci. 2009;106: 15055–15060. doi:10.1073/pnas.0905110106

26. Špaček J. Three-dimensional analysis of dendritic spines. II. Spine apparatus and other cytoplasmic components. Anat Embryol (Berl). 1985;171: 235–243. doi:10.1007/BF00341418

27. Pierce JP, Mayer T, McCarthy JB. Evidence for a satellite secretory pathway in neuronal dendritic spines. Curr Biol. 2001;11: 351–355. doi:10.1016/S0960-9822(01)00077-X

28. Pierce JP, van Leyen K, McCarthy JB. Translocation machinery for synthesis of integral membrane and secretory proteins in dendritic spines. Nat Neurosci. 2000;3: 311–313. doi:10.1038/73868

29. Kruse P, Brandes G, Hemeling H, Huang Z, Wrede C, Hegermann J, et al. Synaptopodin Regulates Denervation-Induced Plasticity at Hippocampal Mossy Fiber Synapses. Cells. 2024;13: 114. doi:10.3390/cells13020114

30. Fukazawa Y, Saitoh Y, Ozawa F, Ohta Y, Mizuno K, Inokuchi K. Hippocampal LTP Is Accompanied by Enhanced F-Actin Content within the Dendritic Spine that Is Essential for Late LTP Maintenance In Vivo. Neuron. 2003;38: 447–460. doi:10.1016/S0896-6273(03)00206-X

31. Korkotian E, Frotscher M, Segal M. Synaptopodin Regulates Spine Plasticity: Mediation by Calcium Stores. J Neurosci. 2014;34: 11641–11651. doi:10.1523/JNEUROSCI.0381-14.2014

32. Vlachos A, Korkotian E, Schonfeld E, Copanaki E, Deller T, Segal M. Synaptopodin Regulates Plasticity of Dendritic Spines in Hippocampal Neurons. J Neurosci. 2009;29: 1017–1033. doi:10.1523/JNEUROSCI.5528-08.2009

33. Chirillo MA, Waters MS, Lindsey LF, Bourne JN, Harris KM. Local resources of polyribosomes and SER promote synapse enlargement and spine clustering after long-term potentiation in adult rat hippocampus. Sci Rep. 2019;9: 3861. doi:10.1038/s41598-019-40520-x

34. Benedeczky I, Molnár E, Somogyi P. The cisternal organelle as a Ca2+-storing compartment associated with GABAergic synapses in the axon initial segment of hippocampal pyramidal neurones. Exp Brain Res. 1994;101: 216–230. doi:10.1007/BF00228742

35. King AN, Manning CF, Trimmer JS. A unique ion channel clustering domain on the axon initial segment of mammalian neurons. J Comp Neurol. 2014;522: 2594–2608. doi:10.1002/cne.23551

36. Lipkin AM, Cunniff MM, Spratt PWE, Lemke SM, Bender KJ. Functional Microstructure of CaV-Mediated Calcium Signaling in the Axon Initial Segment. J Neurosci. 2021;41: 3764–3776. doi:10.1523/JNEUROSCI.2843-20.2021

37. Schlüter A, Del Turco D, Deller T, Gutzmann A, Schultz C, Engelhardt M. Structural Plasticity of Synaptopodin in the Axon Initial Segment during Visual Cortex Development. Cereb Cortex. 2017;27: 4662–4675. doi:10.1093/cercor/bhx208

38. Watson DJ, Ostroff L, Cao G, Parker PH, Smith H, Harris KM. LTP enhances synaptogenesis in the developing hippocampus. Hippocampus. 2016;26: 560–576. doi:10.1002/hipo.22536

39. Harris KM. Structural LTP: from synaptogenesis to regulated synapse enlargement and clustering. Curr Opin Neurobiol. 2020;63: 189–197. doi:10.1016/j.conb.2020.04.009

40. Mundel P, Heid HW, Mundel TM, Kruger M, Reiser J, Kriz W. Synaptopodin: An Actin-associated Protein in Telencephalic Dendrites and Renal Podocytes. J Cell Biol. 1997;139: 193–204. doi:10.1083/jcb.139.1.193

41. Czarnecki K, Haas CA, Bas Orth C, Deller T, Frotscher M. Postnatal development of synaptopodin expression in the rodent hippocampus. J Comp Neurol. 2005;490: 133–144. doi:10.1002/cne.20651

42. Špaček J, Harris KM. Three-Dimensional Organization of Smooth Endoplasmic Reticulum in Hippocampal CA1 Dendrites and Dendritic Spines of the Immature and Mature Rat. J Neurosci. 1997;17: 190–203.

43. Cooney JR, Hurlburt JL, Selig DK, Harris KM, Fiala JC. Endosomal Compartments Serve Multiple Hippocampal Dendritic Spines from a Widespread Rather Than a Local Store of Recycling Membrane. J Neurosci. 2002;22: 2215–2224.

44. Jedlicka P, Schwarzacher SW, Winkels R, Kienzler F, Frotscher M, Bramham CR, et al. Impairment of in vivo theta-burst long-term potentiation and network excitability in the dentate gyrus of synaptopodin-deficient mice lacking the spine apparatus and the cisternal organelle. Hippocampus. 2009;19: 130–140. doi:10.1002/hipo.20489

45. Zhang X, Pöschel B, Faul C, Upreti C, Stanton PK, Mundel P. Essential Role for Synaptopodin in Dendritic Spine Plasticity of the Developing Hippocampus. J Neurosci. 2013;33: 12510–12518. doi:10.1523/JNEUROSCI.2983-12.2013

46. Grigoryan G, Segal M. Ryanodine-mediated conversion of STP to LTP is lacking in synaptopodin-deficient mice. Brain Struct Funct. 2016;221: 2393–2397. doi:10.1007/s00429-015-1026-7

47. Wu PY, Inglebert Y, McKinney RA. Synaptopodin: a key regulator of Hebbian plasticity. Front Cell Neurosci. 2024;18. doi:10.3389/fncel.2024.1482844

48. Inglebert Y, Wu PY, Tourbina-Kolomiets J, Dang CL, McKinney RA. Synaptopodin is required for long-term depression at Schaffer collateral-CA1 synapses. Mol Brain. 2024;17: 17. doi:10.1186/s13041-024-01089-3

49. Thomas KR, Capecchi MR. Site-directed mutagenesis by gene targeting in mouse embryo-derived stem cells. Cell. 1987;51: 503–512. doi:10.1016/0092-8674(87)90646-5

50. Shao Y, Guan Y, Wang L, Qiu Z, Liu M, Chen Y, et al. CRISPR/Cas-mediated genome editing in the rat via direct injection of one-cell embryos. Nat Protoc. 2014;9: 2493–2512. doi:10.1038/nprot.2014.171

51. Ostrovskaya OI, Cao G, Eroglu C, Harris KM. Developmental onset of enduring long-term potentiation in mouse hippocampus. Hippocampus. 2020;30: 1298–1312. doi:10.1002/hipo.23257

52. Kirov SA, Goddard CA, Harris KM. Age-dependence in the homeostatic upregulation of hippocampal dendritic spine number during blocked synaptic transmission. Neuropharmacology. 2004;47: 640–648. doi:10.1016/j.neuropharm.2004.07.039

53. Cao G, Harris KM. Developmental regulation of the late phase of long-term potentiation (L-LTP) and metaplasticity in hippocampal area CA1 of the rat. J Neurophysiol. 2012;107: 902–912. doi:10.1152/jn.00780.2011

54. Hok V, Poucet B, Duvelle É, Save É, Sargolini F. Spatial cognition in mice and rats: similarities and differences in brain and behavior. WIREs Cogn Sci. 2016;7: 406–421. doi:10.1002/wcs.1411

55. Meijer MK, Sommer R, Spruijt BM, van Zutphen LFM, Baumans V. Influence of environmental enrichment and handling on the acute stress response in individually housed mice. Lab Anim. 2007;41: 161–173. doi:10.1258/002367707780378168

56. Ellenbroek B, Youn J. Rodent models in neuroscience research: is it a rat race? Dis Model Mech. 2016;9: 1079– 1087. doi:10.1242/dmm.026120

57. Al Banchaabouchi M, Marescau B, Possemiers I, D’Hooge R, Levillain O, De Deyn PP. NG,NG-Dimethylarginine and NG,N’G-dimethylarginine in renal insufficiency. Pflüg Arch. 2000;439: 524–531. doi:10.1007/s004249900220

58. Coyne MJ, Schultze AE, Iii DJM, Murphy RE, Cross J, Strong-Townsend M, et al. Evaluation of renal injury and function biomarkers, including symmetric dimethylarginine (SDMA), in the rat passive Heymann nephritis (PHN) model. PLOS ONE. 2022;17: e0269085. doi:10.1371/journal.pone.0269085

59. Hamlin DM, Schultze AE, Coyne MJ, McCrann DJI, Mack R, Drake C, et al. Evaluation of Renal Biomarkers, Including Symmetric Dimethylarginine, following Gentamicin-Induced Proximal Tubular Injury in the Rat. Kidney 360. 2022;3: 341. doi:10.34067/KID.0006542020

60. Kohnken R, Himmel L, Logan M, Peterson R, Biswas S, Dunn C, et al. Symmetric Dimethylarginine Is a Sensitive Biomarker of Glomerular Injury in Rats. Toxicol Pathol. 2022;50: 176–185. doi:10.1177/01926233211045341

61. Fiala JC, Harris KM. Extending Unbiased Stereology of Brain Ultrastructure to Three-dimensional Volumes. J Am Med Inform Assoc. 2001;8: 1–16. doi:10.1136/jamia.2001.0080001

62. Palay SL, Sotelo C, Peters A, Orkand PM. THE AXON HILLOCK AND THE INITIAL SEGMENT. J Cell Biol. 1968;38: 193–201. doi:10.1083/jcb.38.1.193

63. Peters A, Proskauer CC, Kaiserman-Abramof IR. THE SMALL PYRAMIDAL NEURON OF THE RAT CEREBRAL CORTEX: The Axon Hillock and Initial Segment. J Cell Biol. 1968;39: 604–619. doi:10.1083/jcb.39.3.604

64. Moor MB, Bonny O. Ways of calcium reabsorption in the kidney. Am J Physiol-Ren Physiol. 2016;310: F1337– F1350. doi:10.1152/ajprenal.00273.2015

65. Staruschenko A, Alexander RT, Caplan MJ, Ilatovskaya DV. Calcium signalling and transport in the kidney. Nat Rev Nephrol. 2024;20: 541–555. doi:10.1038/s41581-024-00835-z

66. Klintsova AY, Cowell RM, Swain RA, Napper RMA, Goodlett CR, Greenough WT. Therapeutic effects of complex motor training on motor performance deficits induced by neonatal binge-like alcohol exposure in rats: I. Behavioral results. Brain Res. 1998;800: 48–61. doi:10.1016/S0006-8993(98)00495-8

67. McFadyen MP, Kusek G, Bolivar VJ, Flaherty L. Differences among eight inbred strains of mice in motor ability and motor learning on a rotorod. Genes Brain Behav. 2003;2: 214–219. doi:10.1034/j.1601-183X.2003.00028.x

68. Tanaka S, Young JW, Halberstadt AL, Masten VL, Geyer MA. Four factors underlying mouse behavior in an open field. Behav Brain Res. 2012;233: 55–61. doi:10.1016/j.bbr.2012.04.045

69. Schultz RL, Karlsson U. Spine apparatus occurrence during different fixation procedures. J Ultrastruct Res. 1966;14: 268–276. doi:10.1016/S0022-5320(66)80048-5

70. Westrum LE, Jones DH, Gray EG, Barron J. Microtubules, dendritic spines and spine apparatuses. Cell Tissue Res. 1980;208: 171–181. doi:10.1007/BF00234868

71. Terasaki M, Reese TS. Characterization of endoplasmic reticulum by co-localization of BiP and dicarbocyanine dyes. J Cell Sci. 1992;101: 315–322. doi:10.1242/jcs.101.2.315

72. Martone M, Zhang Y, Simpliciano V, Carragher B, Ellisman M. Three-dimensional visualization of the smooth endoplasmic reticulum in Purkinje cell dendrites. J Neurosci. 1993;13: 4636–4646.

73. Ehlers MD. Dendritic trafficking for neuronal growth and plasticity. Biochem Soc Trans. 2013;41: 1365–1382. doi:10.1042/BST20130081

74. Martínez G, Khatiwada S, Costa-Mattioli M, Hetz C. ER Proteostasis Control of Neuronal Physiology and Synaptic Function. Trends Neurosci. 2018;41: 610–624. doi:10.1016/j.tins.2018.05.009

75. Perez-Alvarez A, Yin S, Schulze C, Hammer JA, Wagner W, Oertner TG. Endoplasmic reticulum visits highly active spines and prevents runaway potentiation of synapses. Nat Commun. 2020;11: 5083. doi:10.1038/s41467-020-18889-5

76. Deller T, Bas Orth C, Del Turco D, Vlachos A, Burbach GJ, Drakew A, et al. A role for synaptopodin and the spine apparatus in hippocampal synaptic plasticity. Ann Anat - Anat Anz. 2007;189: 5–16. doi:10.1016/j.aanat.2006.06.013

77. Cui-Wang T, Hanus C, Cui T, Helton T, Bourne J, Watson D, et al. Local Zones of Endoplasmic Reticulum Complexity Confine Cargo in Neuronal Dendrites. Cell. 2012;148: 309–321. doi:10.1016/j.cell.2011.11.056

79. Gilbert ET, Klaver LMF, Arndt KC, Kim J, Jia X, McKenzie S, et al. Reciprocal interactions between CA1 pyramidal and axo-axonic cells control sharp wave-ripple events. bioRxiv; 2025. p. 2024.07.02.601726. doi:10.1101/2024.07.02.601726

78. Gallo NB, Paul A, Aelst LV. Shedding Light on Chandelier Cell Development, Connectivity, and Contribution to Neural Disorders. Trends Neurosci. 2020;43: 565–580. doi:10.1016/j.tins.2020.05.003

80. Schneider-Mizell CM, Bodor AL, Collman F, Brittain D, Bleckert A, Dorkenwald S, et al. Structure and function of axo-axonic inhibition. Calabrese RL, Callaway E, Huang ZJ, Oberlaender M, editors. eLife. 2021;10: e73783. doi:10.7554/eLife.73783

81. Dudok B, Szoboszlay M, Paul A, Klein PM, Liao Z, Hwaun E, et al. Recruitment and inhibitory action of hippocampal axo-axonic cells during behavior. Neuron. 2021;109: 3838–3850.e8. doi:10.1016/j.neuron.2021.09.033

82. Aika Y, Ren JQ, Kosaka K, Kosaka T. Quantitative analysis of GABA-like-immunoreactive and parvalbumin- containing neurons in the CA1 region of the rat hippocampus using a stereological method, the disector. Exp Brain Res. 1994;99: 267–276. doi:10.1007/BF00239593

83. Jinno S, Kosaka T. Stereological estimation of numerical densities of glutamatergic principal neurons in the mouse hippocampus. Hippocampus. 2010;20: 829–840. doi:10.1002/hipo.20685

84. Routh BN, Johnston D, Harris K, Chitwood RA. Anatomical and Electrophysiological Comparison of CA1 Pyramidal Neurons of the Rat and Mouse. J Neurophysiol. 2009;102: 2288–2302. doi:10.1152/jn.00082.2009

85. Vitale P, Librizzi F, Vaiana AC, Capuana E, Pezzoli M, Shi Y, et al. Different responses of mice and rats hippocampus CA1 pyramidal neurons to in vitro and in vivo-like inputs. Front Cell Neurosci. 2023;17. doi:10.3389/fncel.2023.1281932

86. Bourne JN, Harris KM. Coordination of size and number of excitatory and inhibitory synapses results in a balanced structural plasticity along mature hippocampal CA1 dendrites during LTP. Hippocampus. 2011;21: 354–373. doi:10.1002/hipo.20768

87. Raymond CR. LTP forms 1, 2 and 3: different mechanisms for the ‘long’ in long-term potentiation. Trends Neurosci. 2007;30: 167–175. doi:10.1016/j.tins.2007.01.007

88. Špaček J. Ultrastructural pathology of dendritic spines in epitumorous human cerebral cortex. Acta Neuropathol (Berl). 1987;73: 77–85. doi:10.1007/BF00695505

89. Reddy PH, Mani G, Park BS, Jacques J, Murdoch G, Whetsell Jr. W, et al. Differential loss of synaptic proteins in Alzheimer’s disease: Implications for synaptic dysfunction. J Alzheimer’s Dis. 2005;7: 103–117. doi:10.3233/JAD-2005-7203

90. Vasil’ev DS, Tumanova NL, Zhuravin IA. [Study of distribution of protein of the spine apparatus synaptopodin in cortical brain parts of rats submitted to hypoxia at different periods of embryogenesis]. Zhurnal Evoliutsionnoĭ Biokhimii Fiziol. 2010;46: 435–439.

91. Cohen JW, Louneva N, Han L, Hodes GE, Wilson RS, Bennett DA, et al. Chronic corticosterone exposure alters postsynaptic protein levels of PSD 95, NR1, and synaptopodin in the mouse brain. Synapse. 2011;65: 763–770. doi:10.1002/syn.20900

92. Arnold SE, Louneva N, Cao K, Wang L-S, Han L-Y, Wolk DA, et al. Cellular, synaptic, and biochemical features of resilient cognition in Alzheimer’s disease. Neurobiol Aging. 2013;34: 157–168. doi:10.1016/j.neurobiolaging.2012.03.004

93. Strehl A, Lenz M, Itsekson-Hayosh Z, Becker D, Chapman J, Deller T, et al. Systemic inflammation is associated with a reduction in Synaptopodin expression in the mouse hippocampus. Exp Neurol. 2014;261: 230–235. doi:10.1016/j.expneurol.2014.04.033

94. Sauer B, Henderson N. Site-specific DNA recombination in mammalian cells by the Cre recombinase of bacteriophage P1. Proc Natl Acad Sci. 1988;85: 5166–5170. doi:10.1073/pnas.85.14.5166

95. Schönig K, Weber T, Frömmig A, Wendler L, Pesold B, Djandji D, et al. Conditional gene expression systems in the transgenic rat brain. BMC Biol. 2012;10: 77. doi:10.1186/1741-7007-10-77

96. Fornasiero EF, Mandad S, Wildhagen H, Alevra M, Rammner B, Keihani S, et al. Precisely measured protein lifetimes in the mouse brain reveal differences across tissues and subcellular fractions. Nat Commun. 2018;9: 4230. doi:10.1038/s41467-018-06519-0

97. Concordet J-P, Haeussler M. CRISPOR: intuitive guide selection for CRISPR/Cas9 genome editing experiments and screens. Nucleic Acids Res. 2018;46: W242–W245. doi:10.1093/nar/gky354

98. Iyer KA, Tenchov R, Lotti Diaz LM, Jain P, Thite T, Deng Y, et al. CRISPR Technology: Transforming the Future of Medicine and Diagnostics. Biochemistry. 2025;64: 4628–4660. doi:10.1021/acs.biochem.5c00480

99. Marcondes FK, Bianchi FJ, Tanno AP. Determination of the estrous cycle phases of rats: some helpful considerations. Braz J Biol. 2002;62: 609–614. doi:10.1590/S1519-69842002000400008

100. Byers SL, Wiles MV, Dunn SL, Taft RA. Mouse Estrous Cycle Identification Tool and Images. PLOS ONE. 2012;7: e35538. doi:10.1371/journal.pone.0035538

101. Becker JB, Arnold AP, Berkley KJ, Blaustein JD, Eckel LA, Hampson E, et al. Strategies and Methods for Research on Sex Differences in Brain and Behavior. Endocrinology. 2005;146: 1650–1673. doi:10.1210/en.2004-1142

102. Paxinos G, Watson C. The rat brain in stereotaxic coordinates. 6th Edition. London: Academic Press; 2007.

103. Schindelin J, Arganda-Carreras I, Frise E, Kaynig V, Longair M, Pietzsch T, et al. Fiji: an open-source platform for biological-image analysis. Nat Methods. 2012;9: 676–682. doi:10.1038/nmeth.2019

104. Kuwajima M, Mendenhall JM, Harris KM. Large-Volume Reconstruction of Brain Tissue from High-Resolution Serial Section Images Acquired by SEM-Based Scanning Transmission Electron Microscopy. Methods Mol Biol Clifton NJ. 2013;950: 253–273. doi:10.1007/978-1-62703-137-0_15

105. Mascorro JA. Novel choices for formulating embedding media kits. Scanning Microscopy 2009. International Society for Optics and Photonics; 2009. p. 73780F. doi:10.1117/12.821821

106. Kuwajima M, Mendenhall JM, Lindsey LF, Harris KM. Automated Transmission-Mode Scanning Electron Microscopy (tSEM) for Large Volume Analysis at Nanoscale Resolution. PLoS ONE. 2013;8: e59573. doi:10.1371/journal.pone.0059573

107. Wetzel AW, Bakal J, Dittrich M, Hildebrand DGC, Morgan JL, Lichtman JW. Registering large volume serial-section electron microscopy image sets for neural circuit reconstruction using FFT signal whitening. 2016 IEEE Applied Imagery Pattern Recognition Workshop (AIPR). 2016. pp. 1–10. doi:10.1109/AIPR.2016.8010595

108. Chirillo MA, Falco JN, Musslewhite MD, Lindsey LF, Harris KM. PyReconstruct: A fully open-source, collaborative successor to Reconstruct. Proc Natl Acad Sci. 2025;122: e2505822122. doi:10.1073/pnas.2505822122

109. Fiala JC, Harris KM. Cylindrical diameters method for calibrating section thickness in serial electron microscopy. J Microsc. 2001;202: 468–472. doi:10.1046/j.1365-2818.2001.00926.x

110. Reymann KG, Frey JU. The late maintenance of hippocampal LTP: Requirements, phases, ‘synaptic tagging’, ‘late-associativity’ and implications. Neuropharmacology. 2007;52: 24–40. doi:10.1016/j.neuropharm.2006.07.026

