## Supplementary material for "*Synaptopodin* KO rat for assessing the dendritic spine apparatus and axonal cisternal organelle in synaptic plasticity, development, and behavior": S1 Document

### S1 Document. Information on CRISPR Reagents and Genotyping.

#### Index

1. CRISPR Reagents ........................................................... Page 1
2. Genotyping Information
   a. LE-*Synpo*$^{em1Kmh}$ .................................................. Page 2
   b. LE-*Synpo*$^{em2Kmh}$ .................................................. Page 4

---

### CRISPR Reagents

#### 5’ guide RNA sequence

`AAAGGTAAACACGTCAGACG`

#### 3’ guide RNA sequence

`GTCAAGGGGCTACTCCAGTA`

---

### Genotyping Information

#### LE-*Synpo*$^{em1Kmh}$ (also known as "synaptopodin knockout rat allele A"; RRID:RRRC_00964; RGD 155782907)

##### *Wildtype*

**5' region sequence** = 939 bp

```
TGGAGTGACTCAGCCTTTGCCACCTGGAGCTGTACGGCCTTGGGTCTGTCAGGAATCTCTGAACTTCTTCACACAG
CTGCCCAGTGGCTGCCTCTCAGTACACTGGGGAGAACAAGATGAATGAACACAGGGAGCGTTGACATTAGCCCAGG
GTATCACTTGGCTTATGTTAGCATGTACAAATTTTTAAATAATTATCTTAACAAAAGTGAGAGGAAAAAAATGGGG
AAGGGTGTAGGAGAAACTGGATCGTGGTCTCTTGATTTGATGTGTTATAATTAGGGTTGGGGACACGGCTACTTTT
TCATGTTTTTATTTTTTAGAACTAAAACGTGTTAACCATTTTATTTTTTGGAGACATTAGTCCTGGTTGTCCTGGA
GCATGTTATGTAGACCGGGCTGGCCTAGAACTTGCAAAGATCCTCCTGCCTCTGCTTTCCCAAGTGCTGGGATTAT
AGGCATGTATCACCATGCCTGCATATTAATATGTTTTCCGAAAAAGAAGTCTTAGGGGCAGGAGAGATGGCTCAAC
AGGTAAAGGTACTTGCTGTGCAAGCCTGGTGTCTGAGTTTGATTCCCAGGACCCGTGTAAAGGTAGAACGAGACAA
AATTATCCTCTGACCTCCACATGTGTAAACATGGTACATGCCTCCCCACATCAATAATGATAAATTTGAAAGGCCT
TAAAAAACAACTTAGTGGCCCAGTTCTAAAATGATATCAAGCAAATGACCAAACTTTGAAAGTATAAAGAGAGAAA
TGCGTAAGGTGTCCTGGGCTTTGCACACTGGGGAATCTTTAGGGAATCTATTCTTGGAATTGAGGTCAGAAAGGTA
AACACGTCAGACGAGGCCTTCGGAGCTGTGCTTGAGGTGAGGAGATGGATGGTCAGTCAGGAGATAAGCAAACTCG
TTGATTTCGGACAGTGGTGAATGAGTG
```

Magenta = PCR primer targets

- 5' Forward primer = TGGAGTGACTCAGCCTTTGC
- 5' Reverse primer = CACTCATTCACCACTGTCCGA

Cyan = guide RNA target

Green = 5' breakpoint

**3' region sequence** = 483 bp

```
TCCCAGACCCATGCTTTTCTGGGATCATTGATGGCTAGTTTTAGGTTTTCAAAACCGGGAAGAGACGAGATGGCAG
TCTAACACGGCTACCCTAGTCTAGGGTGTATTACCTATTTGGAAAGGGGTCCATGTTGCAGAATATGCAAGTCATG
CCCAGGATGCGCGGAGGAGACTCCGGTATTGGGTAGAGTTGACTCCTGACCTCTGAGGATTCTTGAAACTCACTCC
ATAGTATTAGAAAGCAAGGGAGGGTGAAGGGGGAGGTTCACAGGGCTTGAATTCAGAGACAAACATGGGTGGGCAG
TCAAGGGGCTACTCCAGTATGGGGCTGCCATAGTGTCTATATTATGCGCAGAAGAATGGGTATTGTTATATTCATT
GTCTTCACATCTCTCTCTGGCCTGGCTCATACCTGTATCAGCCTTTGAGGCATCTGTCCCCTTAACCTCAGTGACT
CTGATACCAGTCCCCAAGAGTGAGTCG
```

Magenta = PCR primer targets

- 3' Forward primer = TCCCAGACCCATGCTTTTCT
- 3' Reverse primer = CGACTCACTCTTGGGGACTG

Cyan = guide RNA target

Yellow = 3' breakpoint

#### *Knockout*

**Deletion allele sequence** = 966 bp

TGGAGTGACTCAGCCTTTGCCACCTGGAGCTGTACGGCCTTGGGTCTGTCAGGAATCTCTGAACTTCTTCACACAG
CTGCCCAGTGGCTGCCTCTCAGTACACTGGGGAGAACAAGATGAATGAACACAGGGAGCGTTGACATTAGCCCAGG
GTATCACTTGGCTTATGTTAGCATGTACAAATTTTTAAATAATTATCTTAACAAAAGTGAGAGGAAAAAAATGGGG
AAGGGTGTAGGAGAAACTGGATCGTGGTCTCTTGATTTGATGTGTTATAATTAGGGTTGGGGACACGGCTACTTTT
TCATGTTTTTATTTTTTAGAACTAAAACGTGTTAACCATTTTATTTTTTGGAGACATTAGTCCTGGTTGTCCTGGA
GCATGTTATGTAGACCGGGCTGGCCTAGAACTTGCAAAGATCCTCCTGCCTCTGCTTTCCCAAGTGCTGGGATTAT
AGGCATGTATCACCATGCCTGCATATTAATATGTTTTCCGAAAAAGAAGTCTTAGGGGCAGGAGAGATGGCTCAAC
AGGTAAAGGTACTTGCTGTGCAAGCCTGGTGTCTGAGTTTGATTCCCAGGACCCGTGTAAAGGTAGAACGAGACAA
AATTATCCTCTGACCTCCACATGTGTAAACATGGTACATGCCTCCCCACATCAATAATGATAAATTTGAAAGGCCT
TAAAAAACAACTTAGTGGCCCAGTTCTAAAATGATATCAAGCAAATGACCAAACTTTGAAAGTATAAAGAGAGAAA
TGCGTAAGGTGTCCTGGGCTTTGCACACTGGGGAATCTTTAGGGAATGGGGCTGCCATAGTGTCTATATTATGCGC
AGAAGAATGGGTATTGTTATATTCATTGTCTTCACATCTCTCTCTGGCCTGGCTCATACCTGTATCAGCCTTTGAG
GCATCTGTCCCCTTAACCTCAGTGACTCTGATACCAGTCCCCAAGAGTGAGTCG

Magenta = PCR primer targets

- PCR product generated using 5’ Forward and 3’ Reverse primers
- 5’ Forward primer = TGGAGTGACTCAGCCTTTGC
- 3’ Reverse primer = CGACTCACTCTTGGGGACTG

Green and Gray = 5’-3’ fusion

### Genotyping Information

#### LE-*Synpo*$^{em2Kmh}$ (also known as "synaptopodin knockout rat allele B"; RRID: RRRC_01025; RGD 616335891)

##### *Wildtype*

**5' region sequence** = 939 bp

TGGAGTGACTCAGCCTTTGCCACCTGGAGCTGTACGGCCTTGGGTCTGTCAGGAATCTCTGAACTTCTTCACACAG
CTGCCCAGTGGCTGCCTCTCAGTACACTGGGGAGAACAAGATGAATGAACACAGGGAGCGTTGACATTAGCCCAGG
GTATCACTTGGCTTATGTTAGCATGTACAAATTTTTAAATAATTATCTTAACAAAAGTGAGAGGAAAAAAATGGGG
AAGGGTGTAGGAGAAACTGGATCGTGGTCTCTTGATTTGATGTGTTATAATTAGGGTTGGGGACACGGCTACTTTT
TCATGTTTTTATTTTTTAGAACTAAAACGTGTTAACCATTTTATTTTTTGGAGACATTAGTCCTGGTTGTCCTGGA
GCATGTTATGTAGACCGGGCTGGCCTAGAACTTGCAAAGATCCTCCTGCCTCTGCTTTCCCAAGTGCTGGGATTAT
AGGCATGTATCACCATGCCTGCATATTAATATGTTTTCCGAAAAAGAAGTCTTAGGGGCAGGAGAGATGGCTCAAC
AGGTAAAGGTACTTGCTGTGCAAGCCTGGTGTCTGAGTTTGATTCCCAGGACCCGTGTAAAGGTAGAACGAGACAA
AATTATCCTCTGACCTCCACATGTGTAAACATGGTACATGCCTCCCCACATCAATAATGATAAATTTGAAAGGCCT
TAAAAAACAACTTAGTGGCCCAGTTCTAAAATGATATCAAGCAAATGACCAAACTTTGAAAGTATAAAGAGAGAAA
TGCGTAAGGTGTCCTGGGCTTTGCACACTGGGGAATCTTTAGGGAATCTATTCTTGGAATTGAGGTCAGAAAGGTA
AACACGTCAGACGAGGCCTTCGGAGCTGTGCTTGAGGTGAGGAGATGGATGGTCAGTCAGGAGATAAGCAAACTCG
TTGATTTCGGACAGTGGTGAATGAGTG

Magenta = PCR primer targets

- 5' Forward primer = TGGAGTGACTCAGCCTTTGC
- 5' Reverse primer = CACTCATTCACCACTGTCCGA

Cyan = guide RNA target

Black = 5' breakpoint

**3' region sequence** = 483 bp

TCCCAGACCCATGCTTTTCTGGGATCATTGATGGCTAGTTTTAGGTTTTCAAAACCGGGAAGAGACGAGATGGCAG
TCTAACACGGCTACCCTAGTCTAGGGTGTATTACCTATTTGGAAAGGGGTCCATGTTGCAGAATATGCAAGTCATG
CCCAGGATGCGCGGAGGAGACTCCGGTATTGGGTAGAGTTGACTCCTGACCTCTGAGGATTCTTGAAACTCACTCC
ATAGTATTAGAAAGCAAGGGAGGGTGAAGGGGGAGGTTCACAGGGCTTGAATTCAGAGACAAACATGGGTGGGCAG
TCAAGGGGCTACTCCAGTATGGGGCTGCCATAGTGTCTATATTATGCGCAGAAGAATGGGTATTGTTATATTCATT
GTCTTCACATCTCTCTCTGGCCTGGCTCATACCTGTATCAGCCTTTGAGGCATCTGTCCCCTTAACCTCAGTGACT
CTGATACCAGTCCCCAAGAGTGAGTCG

Magenta = PCR primer targets

- 3' Forward primer = TCCCAGACCCATGCTTTTCT
- 3' Reverse primer = CGACTCACTCTTGGGGACTG

Cyan = guide RNA target

Gray = 3' breakpoint

#### *Knockout*

**Deletion allele sequence** = 891 bp

TGGAGTGACTCAGCCTTTGCCACCTGGAGCTGTACGGCCTTGGGTCTGTCAGGAATCTCTGAACTTCTTCACACAG
CTGCCCAGTGGCTGCCTCTCAGTACACTGGGGAGAACAAGATGAATGAACACAGGGAGCGTTGACATTAGCCCAGG
GTATCACTTGGCTTATGTTAGCATGTACAAATTTTTAAATAATTATCTTAACAAAAGTGAGAGGAAAAAAATGGGG
AAGGGTGTAGGAGAAACTGGATCGTGGTCTCTTGATTTGATGTGTTATAATTAGGGTTGGGGACACGGCTACTTTT
TCATGTTTTTATTTTTTAGAACTAAAACGTGTTAACCATTTTATTTTTTGGAGACATTAGTCCTGGTTGTCCTGGA
GCATGTTATGTAGACCGGGCTGGCCTAGAACTTGCAAAGATCCTCCTGCCTCTGCTTTCCCAAGTGCTGGGATTAT
AGGCATGTATCACCATGCCTGCATATTAATATGTTTTCCGAAAAAGAAGTCTTAGGGGCAGGAGAGATGGCTCAAC
AGGTAAAGGTACTTGCTGTGCAAGCCTGGTGTCTGAGTTTGATTCCCAGGACCCGTGTAAAGGTAGAACGAGACAA
AATTATCCTCTGACCTCCACATGTGTAAACATGGTACATGCCTCCCCACATCAATAATGATAAATTTGAAAGGCCT
TAAAAAACAACTTAGTGGCCCAGTTCTAAAATGATATCAAGCAAATGACCAGCTGCCATAGTGTCTATATTATGCG
CAGAAGAATGGGTATTGTTATATTCATTGTCTTCACATCTCTCTCTGGCCTGGCTCATACCTGTATCAGCCTTTGA
GGCATCTGTCCCCTTAACCTCAGTGACTCTGATACCAGTCCCCAAGAGTGAGTCG

Magenta = PCR primer targets

- PCR product generated using 5’ Forward and 3’ Reverse primers
- 5’ Forward primer = TGGAGTGACTCAGCCTTTGC
- 3’ Reverse primer = CGACTCACTCTTGGGGACTG

Black and Gray = 5’-3’ fusion
