## Supplementary material for "*Synaptopodin* KO rat for assessing the dendritic spine apparatus and axonal cisternal organelle in synaptic plasticity, development, and behavior": S1 Figure

**S1 Figure. Original gel image used in Figure 1B.**

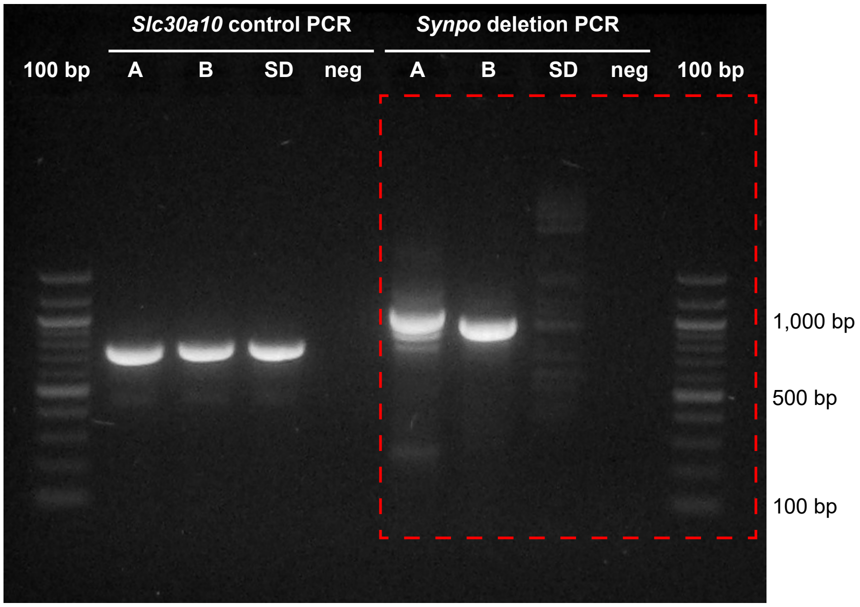


**Left:**
PCR products for *Slc30a10* from two founder lines (A and B), wild-type Sprague-Dawley rat (SD), and negative control (no DNA; neg), showing that DNA in the sample is intact.
The founder lines: A = LE-Synpo$^{em1Kmh}$ and B = LE-Synpo$^{em2Kmh}$

**Right:**
PCR products for *Synpo* from two founder lines show fragments with expected sizes (A, 966 bp; B, 891 bp). For SD, PCR band is absent because the conditions do not support amplification of the longer wild-type sequence. neg = no DNA negative control.
Area indicated by the dotted red box was used in Figure 1B.
